# Brain Networks Detectable by fMRI during On-Line Self Report of Hallucinations in Schizophrenia

**DOI:** 10.1101/2021.11.06.467564

**Authors:** Karanvir Gill, Chantal Percival, Meighen Roes, Leo Arreaza, Abhijit Chinchani, Nicole Sanford, Walter Sena, Homa Mohammadsadeghi, Mahesh Menon, Matthew Hughes, Sean Carruthers, Philip Sumner, Will Woods, Renaud Jardri, Iris E. Sommer, Susan L. Rossell, Todd S. Woodward

## Abstract

An analysis of an internationally shared functional magnetic resonance imaging (fMRI) data involving healthy participants and schizophrenia patients extracted brain networks involved in listening to radio speech and capture hallucination experiences. A multidimensional analysis technique demonstrated that for radio-speech sound files, a brain network matching known auditory perception networks emerged, and importantly, displayed speech-duration-dependent hemodynamic responses (HDRs), confirming fMRI detection of these speech events. In the hallucination-capture data, although a sensorimotor (response) network emerged, it did not show hallucination-duration-dependent HDRs. We conclude that although fMRI retrieved the brain network involved in generating the motor responses indicating the start and end of an experienced hallucination, the hallucination event itself was not detected. Previous reports on brain networks detected by fMRI during hallucination capture is reviewed in this context.

## 1. Introduction

Auditory verbal hallucinations (AVHs) are speech perceptions that occur in the absence of an external stimulus and are primary symptoms of psychosis, such that 60-80% of people with schizophrenia spectrum disorder report experiencing them ^[1, 2]^. The propensity to hallucinate (e.g., experiencing hallucinations in the past week) has been linked with neural hyperactivity in voice-selective regions of the STG ^[3-7]^, and these voice-selective cortical regions have been reported to activate in symptom capture studies on hallucinations ^[8, 9]^. Repeated transcranial magnetic stimulation (rTMS) of voice-selective cortical regions was also reported to reduce the intensity of hallucinations ^[10-12]^. Therefore, much of the AVH literature to date has taken a region-of-interest based approach focusing on voice-selective regions of the STG.

AVHs are unlikely to arise exclusively from hyperactivity in the STG. For example, the breakaway speech/unbidden thoughts account of hallucinations ^[13-15]^ puts forward that AVH may occur when self-monitoring breaks down, possibly due to reduced activation in the dorsal anterior cingulate cortex (dACC)/supplementary motor areas (SMA) during the hallucinatory experience ^[16]^. Metacognitive or belief-based influences are also likely play a role ^[17-19]^; therefore, a network-based approach is important for investigating the biological underpinnings of hallucinations.

In functional magnetic resonance imaging (fMRI) symptom capture studies of hallucinations, activity in the STG is typically reported, as are a number of other language based regions, including Broca’s area, anterior insula, precentral gyrus, frontal operculum, inferior parietal lobule, hippocampus, parahippocampal regions and in motor areas such as the inferior frontal gyrus, cerebellum, insula, and postcentral gyrus ^[20-22]^. In most symptom capture studies, the experimental procedure to monitor hallucinations in the scanner consists of participants pressing a button or squeezing a ball, to indicate the onset and offset of hallucinations ^[20-22]^, with exceptions being relatively rare ^[e.g., 23, 24, 25]^. Since fMRI measures the blood-oxygen-level-dependent (BOLD) signal increases in response to cognitive events, the timing of hallucinations onset/offset is confounded with response, leading to complications interpreting the confounding influence of the sensorimotor (response) network with a potential network underlying hallucination.

In addition to this confound with response processes, another source of potential error in symptom capture studies includes the duration of the hallucinations, which range from one- or two-word utterances to much longer commentary ^[26]^. This, combined with differences between individuals with respect to the accuracy with which they report the duration of their hallucinations, can add noise to the data. An additional source of error affecting fMRI is the dependence on beta weights derived from regressing the BOLD signal onto a synthetic model of the hemodynamic response (HDR) shape ^[27]^. Even if this synthetic HDR shape is adjusted for reported duration of hallucination events, restricting interpreted results exclusively to BOLD signal changes that conform to a model of the HDR will conflate multiple cognitive operations, such as the response indicating hallucination onset/offset with other cognitive processes taking place during hallucinations ^[28]^. This is because the HDR resulting from both the hallucination experience and the response process could partially match the synthetic HDR shape model, making temporal specificity of the involved cognitive processes challenging to retrieve.

In the current study, we used constrained principal component analysis for fMRI ^[fMRI-CPCA; 29, 30]^, which provided dimensional representation of brain networks, whereby retrieved networks can be anatomically compared to previously derived networks which have known functions, such as response and auditory perception ^[31, 32]^. fMRI-CPCA also provides condition- and subject-specific data-driven HDR shapes for each brain network, allowing direct observation of the duration of detected cognitive events for each network separately, without requiring the assumptions/models of the HDR shape that are typically used, as described above. To provide evidence that hallucination-driven HDR have been recorded by fMRI, the following validity requirements needed to be met: (1) spatial validity, (2) temporal validity, and (3) experimental validity ^[33]^. Spatial validity requires observation of known network configurations ^[31, 32, 34-36]^. For example, the sensorimotor (response) network is expected to involve brain regions such as the bilateral supplementary motor area (SMA), dorsal anterior cingulate cortex (dACC), and insula, as well as left somatomotor areas and right cerebellum for a one-handed response ^[30, Figure 7 and Table 6, 31]^, but the auditory perception network is expected to be dominated by the superior temporal gyrus ^[31, 37, Component 7, Figure S3]^. For temporal validity, a biologically plausible HDR shape must be associated with an anatomically valid network to ensure that BOLD signal is likely being detected. For experimental validity, the nature of the HDR shape should change as important experimental conditions and/or measured cognitive operations change; in the case of the current study, the HDR should change with the duration of the radio speech event or experienced hallucination.

Symptom capture data was analyzed by merging separate datasets from two sites (Melbourne and Utrecht), and radio speech events were collected at the Melbourne site only. Our approach was to test spatial, temporal and experimental validity in the external (radio) sound timing from the Melbourne data (i.e., short/medium/long durations for experimental validity), and for the hallucinatory experiences in a data set merged from the two sites (i.e., short/long durations for experimental validity). Temporally, we expect to see a pattern of increased HDR from baseline to peak, with the HDR duration at peak level determined by duration of radio speech/hallucination, before returning back to baseline. By utilizing data from two sites we are able to increase sample sizes so that it was possible to collect together a greater range in both frequency and duration of hallucinations, facilitating identification of these networks.

## 2. Methods

### 2.1 Participants

#### 2.1.1 Melbourne

Seventeen schizophrenia patients and thirty-one healthy control participants were included in the analysis of the radio speech stimuli, and twelve of those schizophrenia patients also contributed data to the symptom capture study completed in Melbourne. Table 1 provides the demographic information of the participants and the scores on the Positive and Negative Syndrome Scale (PANSS) for the schizophrenia patients. The mean age for schizophrenia patients from the external radio voice analysis was 40.06 (SD = 11.87), and for healthy controls it was 31.65 (SD = 11.57). Years of education completed for schizophrenia patients was 14.69 (SD = 2.47), for healthy controls it was 15.90 (SD = 2.15). The mean total PANSS score for schizophrenia patients was 68.53 (SD = 18.44).

**Table 1.** Demographic information of participants and the total PANSS score for the schizophrenia patients from the Melbourne site and psychotic patients from the Utrecht site.

|  | Melbourne |  |  |  |  |  | Utrecht |  |
| --- | --- | --- | --- | --- | --- | --- | --- | --- |
|  | Schizophrenia Patients (radio; n = 17) |  | Healthy Controls (radio; n = 31) |  | Schizophrenia Patients (voices; n = 12) |  | Schizophrenia Patients (n = 15) |  |
|  | N | Mean (SD) | N | Mean (SD) | N | Mean (SD) | N | Mean (SD) |
| **Age** | 17 | 40.06 (11.87)** | 31 | 31.65 (11.57)** | 12 | 42.58 (10.77) | 14* | 37.09 (11.48) |
| **Sex (male/female)** | 7/10 | - | 13/18 | - | 5/7 | - | 6/9 | - |
| **Handedness (right/non-right)** | 12/3* | - | 25/2* | - | 10/1* | - | 8/4* | - |
| **Years of education** | 16* | 14.69 (2.47) | 31 | 15.90 (2.15) | 12 | 14.75 (2.77) | * | - |
| **Total PANSS** | 17 | 68.53 (18.44) | - | - | 12 | 60.50 (11.21) | 10* | 57.53 (15.53) |
| **Positive PANSS** | 17 | 18.76 (5.40) | - | - | 12 | 16.33 (3.47) | 10* | 14.50 (4.03) |
| **PANSS P3 - Hallucinations** | 17 | 4.24 (1.55) | - | - | 12 | 3.83 (1.46) | 10* | 4.80 (.63) |
| **Negative PANSS** | 17 | 15.59 (6.02) | - | - | 12 | 13.58 (4.33) | 10* | 14.0 (4.67) |
| **General Psychopathology** | 17 | 34.18 (9.85) | - | - | 12 | 30.58 (8.14) | 10* | 28.80 (7.87) |

*Note.* \*Information was not available for some participants; \*\*=Melbourne control vs Melbourne schizophrenia patients (n=17), $p < .05$ ; \*\*\*=Melbourne control vs Melbourne schizophrenia patients (n=12), $p < .01$ .

#### 2.1.2 Utrecht

Fifteen schizophrenia patients were included in the analysis of the hallucinations from the symptom capture study completed in Utrecht. Table 1 provides the demographic information of the participants and their scores on the Positive and Negative Syndrome Scale (PANSS). The mean age for the schizophrenia patients was 37.09 (SD = 11.48). The mean total PANSS score for the schizophrenia patients was 57.53 (SD = 15.53).

### 2.2 Tasks

#### 2.2.1 Melbourne

The radio speech task completed by participants from the Melbourne site involved indicating the start and end of (i) radio speech clips, and (ii) experienced hallucinations, using a dominant hand button-press response. All participants participated in a single fMRI acquisition run, and each run consisted of five radio speech blocks and five hallucinations/random pressing blocks (for patients/controls, respectively), eyes closed. These blocks were alternated, one after the other, for the entire run. The random pressing blocks were not analyzed here. A series of on/off tones indicated the change of a block, so that participants with their eyes closed would know when to open them to read the change of instructions, and then were encouraged to close their eyes. During the radio speech block, both schizophrenia patients and control participants pressed one button to indicate the start, and a different button to indicate the end, of an audio clip of radio speech of varying durations. The radio clips were a mixture of male and female voices either independently, or combinations of talking to each other. The tone and content used was neutral, so as to not induce an emotional reaction in participants. During the hallucinations blocks, the schizophrenia patients pressed a button to indicate the start, and another button to indicate end of AVHs.

#### 2.2.2 Utrecht

The task completed by participants from the Utrecht site was to indicate the perception of hallucinations using a balloon squeeze response by their dominant hand. Each subject participated in up to 3 fMRI acquisition runs. During the fMRI acquisition runs, patients indicated the onset of an AVH by a balloon-squeeze, followed by a balloon-release to indicate its offset.

### 2.3 Image Acquisition

Table S1 provides details on the fMRI parameters from both sites.

#### 2.3.1 Melbourne

A Siemens Tim Trio 3 Tesla MRI scanner with an echo planar imaging (EPI) sequence was used, with a repetition time (TR) of 1.150s. The following were the parameter settings: transverse slices, TE = 30ms, flip angle 84°, field of view (FOV) 224 × 224 mm, voxel size 4.5 × 4.5 × 4.5mm with gap = 0.5mm.

#### 2.3.2 Utrecht

A Philips Achieva 3 Tesla MRI scanner with a 3D-PRESTO pulse sequence was used, with a repetition time (TR) of 0.609s. The following were the parameter settings: 40 (coronal) slices, TR/TE 21.75/32.4 ms, flip angle 10°, field of view (FOV) 224 × 256 × 160, matrix 64 × 64 × 40, voxel size 4 mm isotropic.

#### 2.3.3 Preprocessing

Functional scans from both sites (Melbourne and Utrecht) were reoriented to set the origin at the anterior commissure, and the scan series was realigned, coregistered to the accompanying T1 anatomical scan, and normalized using the method implemented in Statistical Parametric Mapping 12 (SPM12; http://www.fil.ion.ucl.ac.uk/spm). Slice-timing correction was applied to the functional scans from the Melbourne dataset because the images were collected using an EPI-sequence, which is not necessary for images collected with PRESTO. All images were normalized by first warping high-resolution structural images to a template of Montreal Neurological Institute (MNI) coordinate space, then applying these transformation parameters to the realigned functional images. Voxels were normalized to 3 × 3 × 3mm. The normalized functional images were smoothed with a Gaussian kernel (8mm FWHM). All X Y Z coordinates listed in this manuscript are MNI coordinates.

From the Melbourne data, if scans from a particular subject revealed translation and/or rotation corrections for head movement exceeding 4mm or 4°, then those participants were excluded from any further analyses. However, from the Utrecht data, since the translation in the y-direction exceeded 4mm, for all participants, in every run, in a similar pattern over the duration of the eight hundred scans, no participants were excluded by this criterion. fMRI-CPCA extracts for analysis only variance predictable from event timing (with hallucinations being the events); therefore, the gradual movement patterns over the entire run is not predictable from event timing, and would be ignored, therefore not affecting the results.

### 2.4 Timing

The sections below outline the details of the various fMRI-CPCA and repeated measures ANOVA analyses conducted with different samples of participants from both sites. Table 2 provides a summary of the number of participants analyzed, tasks, experiment design, mean frequency of hallucinations in participants with schizophrenia per run, mean duration of hallucinations in participants with schizophrenia, and the durations assigned short/medium/long for the analysis of participants from each site. Duration intervals were computed such that there would be an approximately equal number of timing events for each condition. Since the HDR shape is sluggish, taking place over a period of 5-10 seconds, and cannot convolve multiple rapid hallucination events (as indicated by button press or squeeze response), multiple hallucinations of were combined into a single hallucination if the interval between the motor response indicating the terminating a hallucination event, and that indicating the start of another hallucination event, was 3 seconds or less. Table 3 depicts the average frequency of external radio and hallucination responses analyzed per run from both sites, along with the average number of times multiple hallucination responses were combined to be analyzed as a single hallucination, averaged over all subjects.

**Table 2.**
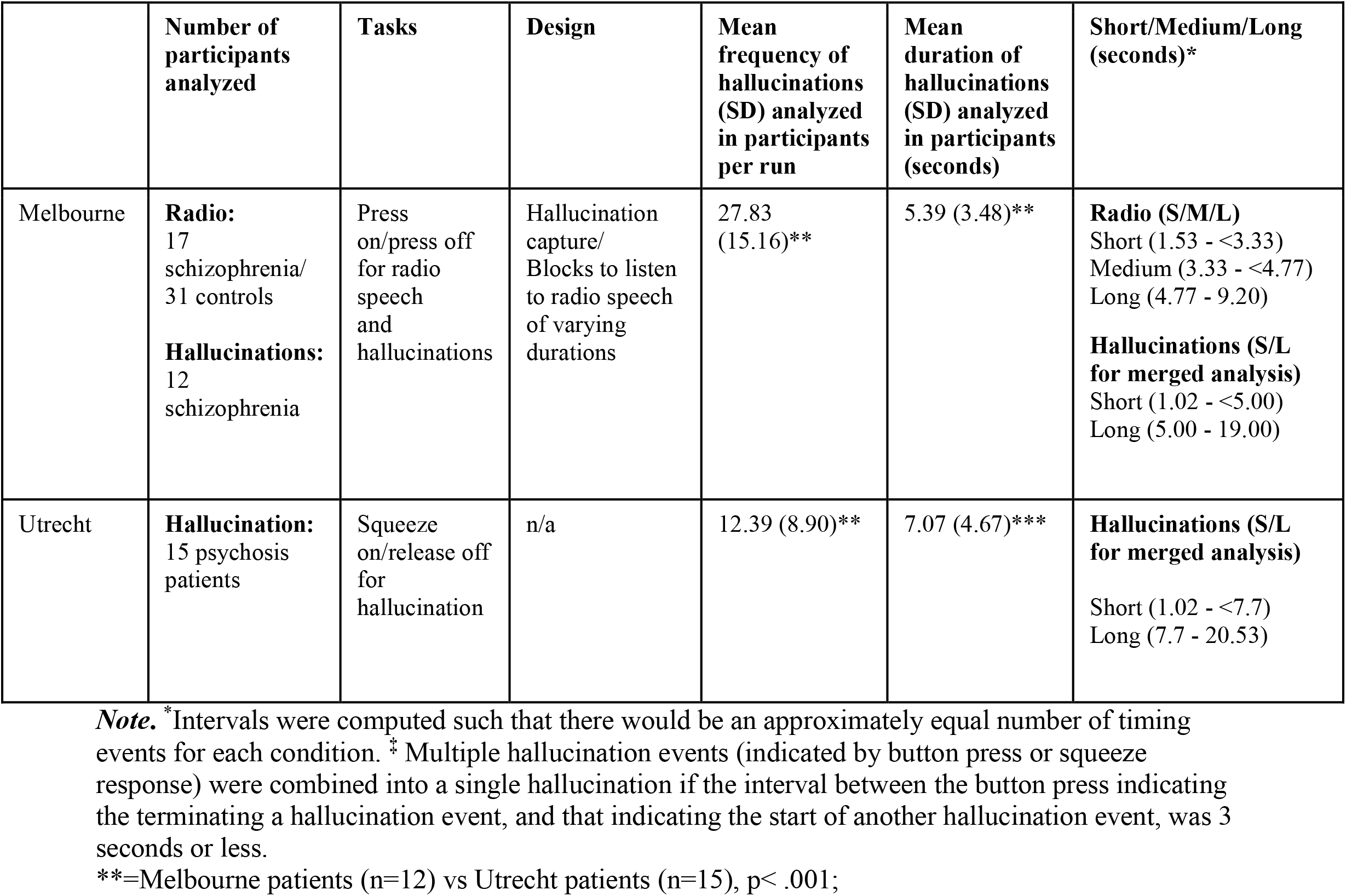
Details of hallucination capture studies acquired from sites in Melbourne and Utrecht

|  | **Number of participants analyzed** | **Tasks** | **Design** | **Mean frequency of hallucinations (SD) analyzed in participants per run** | **Mean duration of hallucinations (SD) analyzed in participants (seconds)** | **Short/Medium/Long (seconds)\*** |
| --- | --- | --- | --- | --- | --- | --- |
| Melbourne | **Radio:**<br>17 schizophrenia/<br>31 controls<br>**Hallucinations:**<br>12 schizophrenia | Press on/press off for radio speech and hallucinations | Hallucination capture/<br>Blocks to listen to radio speech of varying durations | 27.83<br>(15.16)** | 5.39 (3.48)** | **Radio (S/M/L)**<br>Short (1.53 - <3.33)<br>Medium (3.33 - <4.77)<br>Long (4.77 - 9.20)<br>**Hallucinations (S/L for merged analysis)**<br>Short (1.02 - <5.00)<br>Long (5.00 - 19.00) |
| Utrecht | **Hallucination:**<br>15 psychosis patients | Squeeze on/release off for hallucination | n/a | 12.39 (8.90)** | 7.07 (4.67)*** | **Hallucinations (S/L for merged analysis)**<br>Short (1.02 - <7.7)<br>Long (7.7 - 20.53) |

**Note.** \*Intervals were computed such that there would be an approximately equal number of timing events for each condition. † Multiple hallucination events (indicated by button press or squeeze response) were combined into a single hallucination if the interval between the button press indicating the terminating a hallucination event, and that indicating the start of another hallucination event, was 3 seconds or less.
\*\*=Melbourne patients (n=12) vs Utrecht patients (n=15), $p < .001$ ;

**Table 3.** Details of external voice clips and combined hallucinations analyzed in short, medium, and long (S/M/L) intervals.

|  | Melbourne radio (S/M/L) | Melbourne hallucinations (S/M/L; Supplementary Material) |  |  | Melbourne hallucinations (S/L) |  |  | Utrecht hallucinations (S/L) |  |  |
| --- | --- | --- | --- | --- | --- | --- | --- | --- | --- | --- |
|  | Mean frequency of radio clips per run (SD) | Mean frequency of hallucinations analyzed per run (SD) | Mean frequency of hallucinations combined per run (SD) |  | Mean frequency of hallucinations analyzed per run (SD) | Mean frequency of hallucinations combined per run (SD) |  | Mean frequency of hallucinations analyzed per run (SD) | Mean frequency of hallucinations combined per run (SD) |  |
|  |  |  | 2 combined | 3 or more combined |  | 2 combined | 3 or more combined |  | 2 combined | 3 or more combined |
| **Short** | 10.94<br>(0.42) | 9.17<br>(6.01) | 0.50<br>(1.12) | 0.00 | 16.67<br>(10.35) | 0.50<br>(1.12) | 0.00 | 7.57<br>(8.26) | 0.64<br>(1.29) | 0.14<br>(0.58) |
| **Medium** | 12.02<br>(0.80) | 9.42<br>(11.23) | 0.08<br>(0.28) | 0.00 | - | - | - | - | - | - |
| **Long** | 11.76<br>(0.89) | 9.25<br>(6.99) | 0.92<br>(0.95) | 1.00<br>(2.12) | 11.17<br>(8.20) | 1.00<br>(1.00) | 1.00<br>(2.12) | 4.71<br>(2.67) | 0.57<br>(0.98) | 0.57<br>(1.08) |

#### 2.4.1 Melbourne patient/control radio speech (Short/Medium/Long)

The purpose of the Melbourne patient and control radio speech analysis was to provide an example of spatial, temporal and experimental validity, such that if radio speech perception were driving the BOLD signal, then the HDR plot for the brain networks involved in response to auditory stimuli would show staggered HDR peaks for short/medium/long durations, with similar trends as they rise from baseline, but with the three curves peaking at staggered time points (as determined by duration of the radio speech), before returning to baseline.

For the Melbourne patient and control radio speech analysis, timing vectors for the onset of radio speech clips were divided into Short (1.53 - <3.33), Medium (3.33 - <4.77), and Long (4.77 - 9.20), based on radio speech durations (breakdown of frequency of events per duration can be seen in Table 3). BOLD response for full-brain scans 1-20 following radio speech clip onset were estimated using a FIR basis function (i.e., 23 seconds of post-stimulus time with TR = 1,150ms), to allow sufficient time for peak and relaxation of the HDR. Subsequently, within-subjects factors of radio speech duration (Short/Medium/Long) and time (20 post-stimulus time bins) were examined in the resulting predictor weights (which form the HDR), resulting in a 3 (Duration) × 20 (Time) × 2 (Group) ANOVA for each component. Significant effects of Duration × Time, and any effects/interactions involving group differences were further examined. Post hoc analyses of effects involving Duration were carried out using polynomial contrasts (linear *vs*. quadratic trends), and interactions involving Time were broken down using repeated contrasts, which contrast adjacent time bins and adjacent durations as 2 × 2 interactions.

#### 2.4.2 Melbourne and Utrecht patient hallucinations merged (Short/Long)

For the Melbourne and Utrecht patient hallucination analysis, participants from the Melbourne and Utrecht datasets were merged (analysis of each site separately is presented in the Supplementary Material Figures S5-S10). For the Melbourne dataset, timing vectors for the onset of hallucination events in patients were divided into Short (1.02s to <5s) and Long (5s to 19s), with the breakdown of frequency of events per duration presented in Table 3 applied to the 12 patients from the Melbourne dataset with a minimum of 4 hallucination events. Similarly, for the Utrecht dataset, timing vectors for the onset of hallucination events in patients were divided into Short (1.02s to <7.7s) and Long (7.7s to 20.53s), with the breakdown of frequency of events per duration presented in Table 3, applied to the 15 patients with a minimum of 4 hallucination events. The conditions for each dataset were set so that there would be nearly equal number of hallucinations in both short and long categories.

Because the fMRI-CPCA software is currently restricted to an equivalent number of time bins for each group in a merged analysis, we set the number of time bins to scans 1-37 following hallucination onset, as indicated by button press/squeeze response, using a FIR basis function to monitor BOLD response. This resulted in 22.6 seconds of post-stimulus time with TR = 610ms for the Utrecht data, and 42.6 seconds of post-stimulus time with TR = 1,150ms for Melbourne participants. After the regression and PCA steps, inspection of the scree plot of singular values indicated 3 components should be retained. Then, within subject factors of hallucination duration (short vs. long) and time (37 post-stimulus time bins) were examined with the resulting predictor weights, resulting in a 2 (Duration) × 37 (Time) ANOVA for each component. Interactions involving Time were broken down using repeated contrasts, which contrast adjacent time bins and adjacent durations (only two levels in this case) as 2 × 2 interactions.

### 2.5 Data Analysis

Data Analysis was carried out using fMRI-CPCA ^[29, 30]^, as described in detail in the Supplementary Material (also see Figure S1). Table 4 provides assessments of spatial, temporal and experimental validity, as a function of all retrieved networks.

**Table 4.**
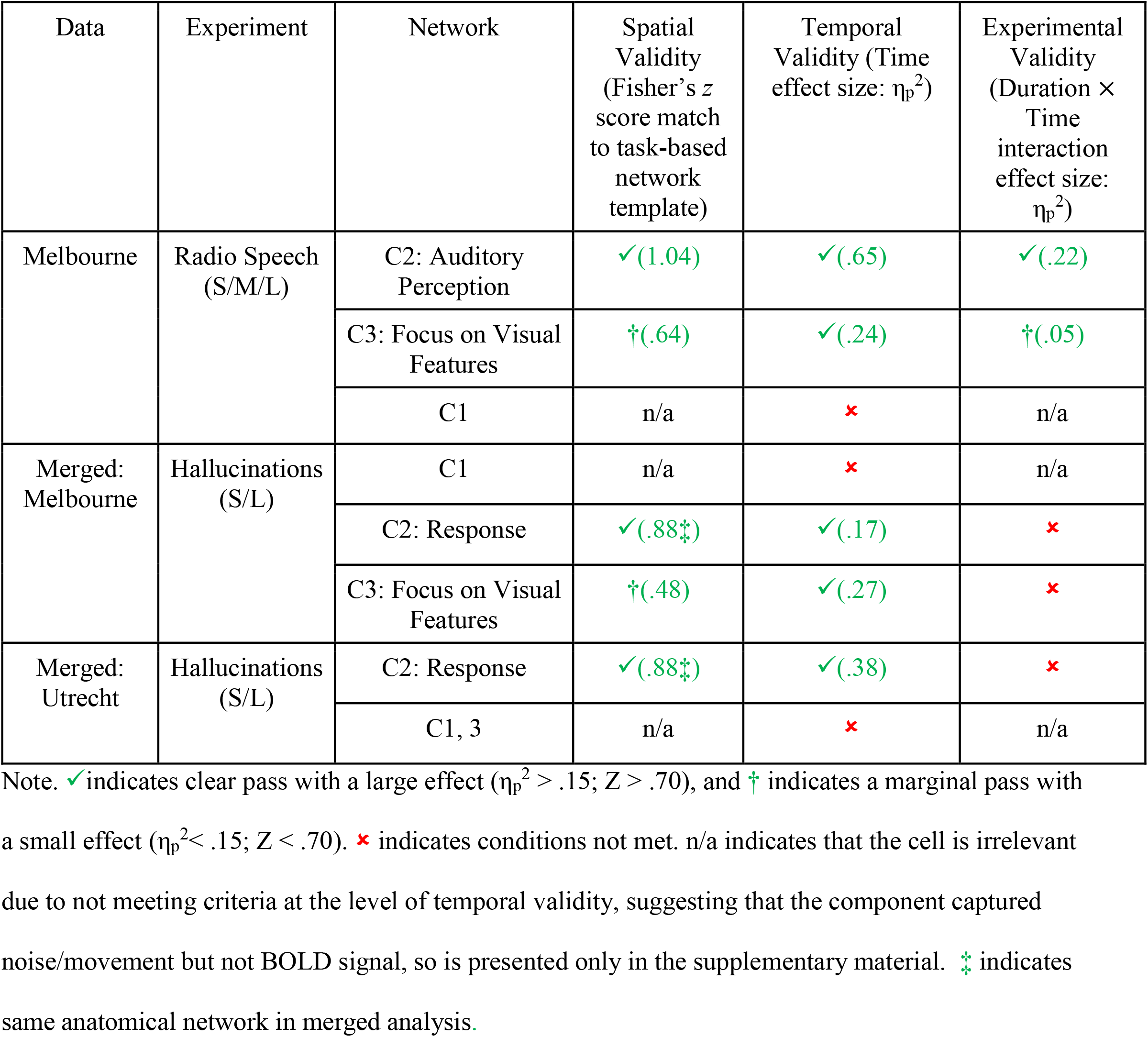
Assessments of Spatial, Temporal and Experimental Validity, as a function of retrieved networks. Fisher-transformed Z scores (in brackets) are reported for spatial validity. Effect sizes (in brackets) are reported for Temporal Validity (detection of biologically plausible HDR shape) are derived from the Time main effect, and those for Experimental Validity (detection of short-medium-long [S/M/L] or short-long [S/L] duration differences in HDR shapes for radio speech or hallucinations event) are derived from the Duration main effect, or Duration ×Time interaction, whichever was largest.

## 3. Results

### 3.1 Melbourne Radio Speech (Short/Medium/Long)

Three components were extracted for the Melbourne radio speech experiment, as determined by examining the scree plot ^[38, 39]^. Component 1 did not retrieve a biologically plausible HDR shape, so is reported only in the supplementary material (Figure S2A/B).

#### 3.1.1 Component 2: Auditory Perception Network

The anatomical regions associated with Component 2 are outlined in Figure 1A and the anatomical description of component two is presented in Table S2. The anatomical pattern matched well to that in the auditory perception network ^[Fisher’s z = 1.04; 31, AUD, 37, Component 7, Figure S3]^, with bilateral peaks in right superior temporal gyrus (xyz: 60, -16, -2), and left planum temporale (xyz: -57 -19, 1).

**Figure 1:**
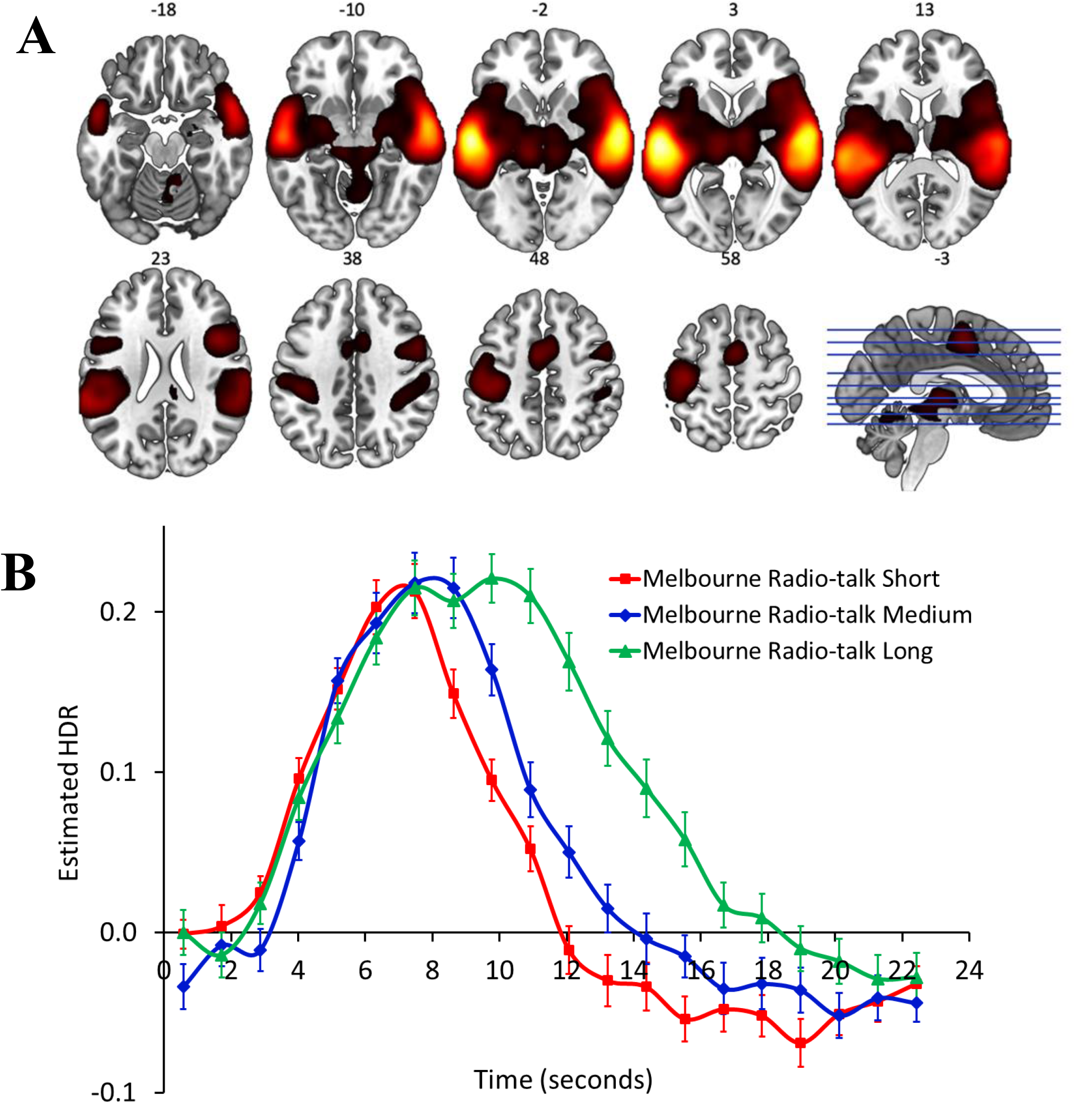
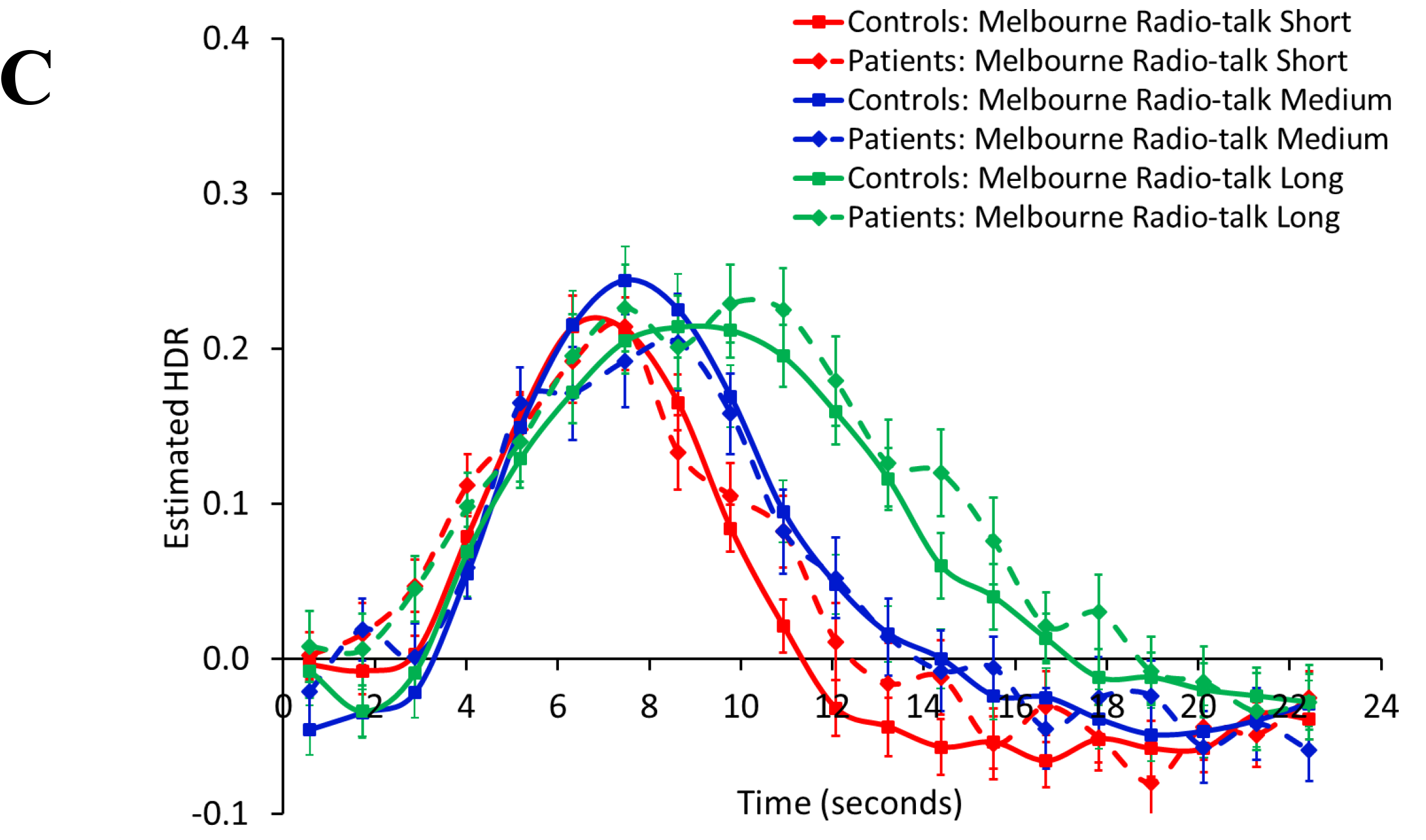
Auditory Perception network, from the Melbourne patient radio speech (Short/Medium/Long) analysis. A (top): dominant 20% of component loadings for Component 2, proposed Auditory Perception network, from the Melbourne patient radio speech (Short/Medium/Long) analysis. MNI Z-axis coordinates are displayed. Images are displayed in neurological convention (left is left). Red/yellow = positive loadings, positive threshold = 0.11, max = 0.41. B (middle): mean finite impulse response (FIR)-based predictor weights plotted as a function of post-stimulus time (TR = 1150ms) and condition (averaged over participants, error bars are standard errors). C (bottom): mean finite impulse response (FIR)-based predictor weights plotted as a function of post-stimulus time (TR = 1150ms) and condition (averaged over participants, error bars are standard errors) shown with group differences.

Figure 1B displays the estimated HDR shape for Component 2. Component 2 displayed a biologically plausible HDR, and a highly significant main effects Time, *F*(19, 874) = 84.31, *p* < 0.001, η_p_^2^ = .65, clearly meeting the requirement of temporal validity. There was also a highly significant main effect of Duration, *F*(2, 92) = 12.45, *p* < 0.001, η_p_^2^ = .21, and an equally strong Duration ×Time interaction, *F*(38, 1748) = 13.02, p < 0.001, η_p_^2^ = .22, which was dominated by differences between Short and Medium duration for the increase from time bins 7 to 8, *F*(1, 46) = 14.08 p < 0.001, η_p_^2^ = .12, differences between Medium and Long for the increase from time bins 4 to 5, *F*(1, 46) = 15.10, *p* < 0.001, η_p_^2^ = .12, the decreases from time bins 8 and 9, and 9 to 10, *F*(1, 46) = 15.05 p < 0.001, η_p_^2^ = .12, *F*(1, 46) = 23.40, *p* < 0.001, η_p_^2^ = .12, respectively. These effects were caused by staggered peaks and increasing durations of activation for the Short, Medium and Long radio speech conditions, respectively, clearly meeting the requirement of experimental validity. This provides strong support for a functional brain network reliably responding to radio speech, displaying strong spatial, temporal and experimental validity ^[33]^ (see Table 4). No main effects or interactions involving Group were significant (all *p*s > .35; see Figure 1C).

#### 3.1.2 Component 3: Focus on Visual Features

The anatomical regions associated with Component 3 are outlined in Figure 2A, and the anatomical description is presented in Table S3. The negative loadings provide a weak match to the Focus on Visual Features (FVF) network (Fisher’s *z* = .64), providing weak evidence for spatial validity. Component 3 is characterized by bilateral deactivation in occipital areas such as the occipital pole (xyz: -27, -91, 16; 15, -88, 28). The FVF network is known to deactivate when the visual details of the task are not relevant to response _[31, FVF, 37, Figure S2, 40, Figure 5.60]_.

**Figure 2:**
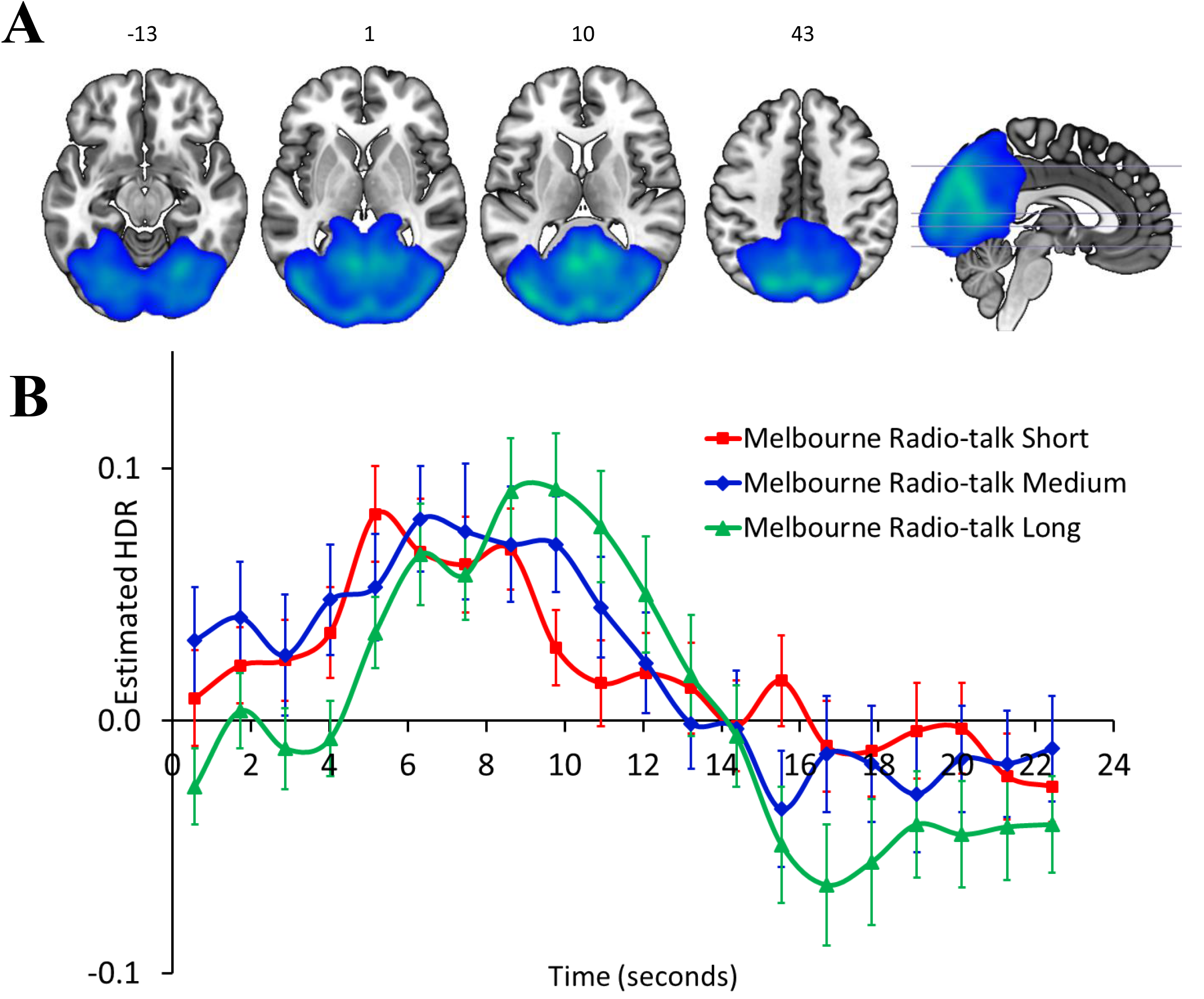
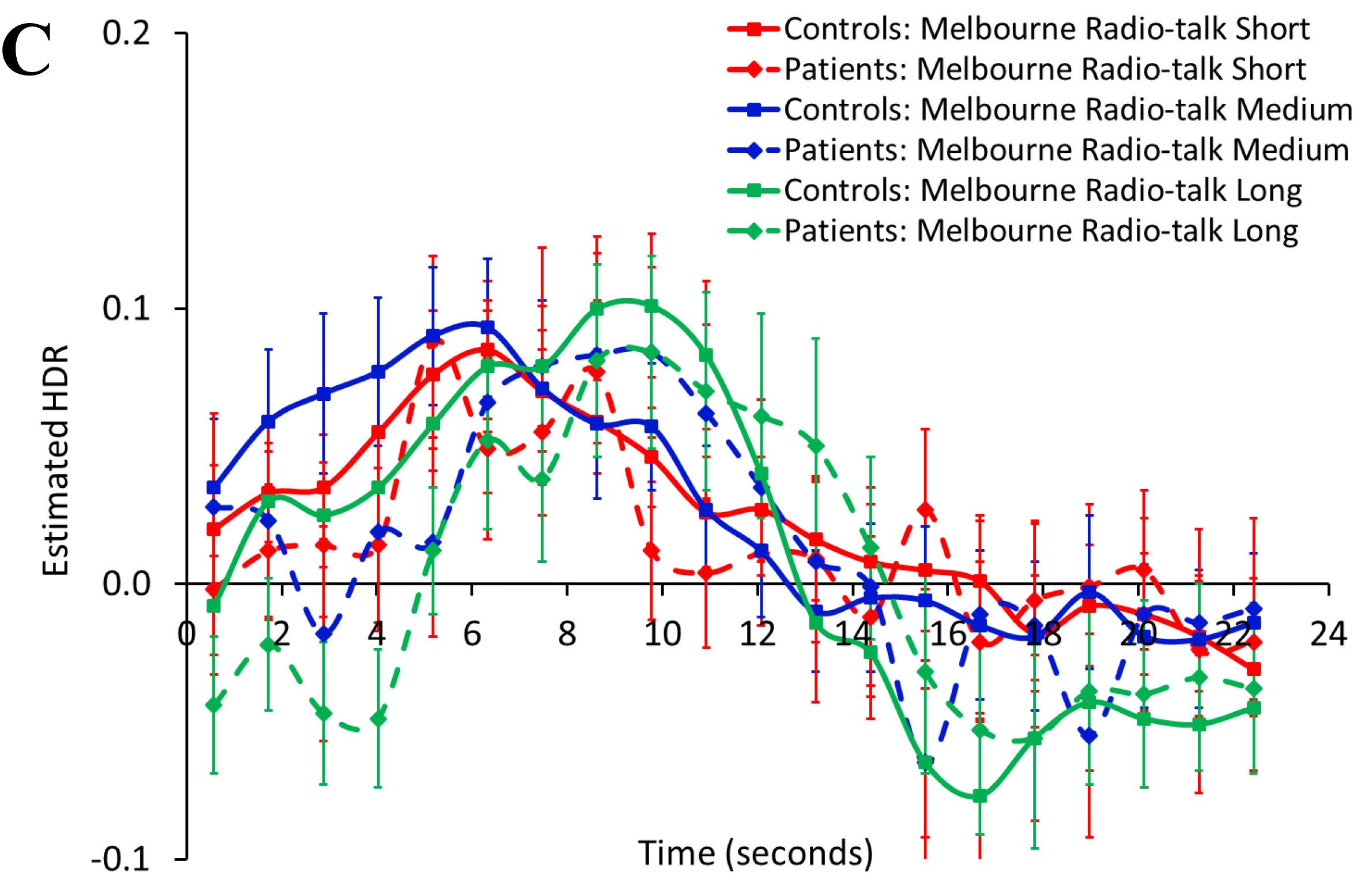
Focus on Visual Features/Auditory Perception network, from the Melbourne radio speech (Short/Medium/Long) analysis. A (top): dominant 20% negative component loadings for Component 3, from the Melbourne radio speech (Short/Medium/Long) analysis, Focus on Visual Features/Auditory Perception. MNI Z-axis coordinates are displayed. Images are displayed in neurological convention (left is left). Blue/green = negative loadings, threshold = -0.12, min = -0.19. B (middle): mean finite impulse response (FIR)-based predictor weights plotted as a function of post-stimulus time (TR = 1150ms) and condition (averaged over participants, error bars are standard errors). C (bottom): mean finite impulse response (FIR)-based predictor weights plotted as a function of post-stimulus time (TR = 1150ms) and condition (averaged over participants, error bars are standard errors) shown with group differences.

Figure 2B displays the estimated HDR shape for Component 3. Component 3 had a significant effect of Time, *F*(19, 874) = 14.70, *p* < 0.001, η_p_^2^ = .24, and although the HDR was biologically plausible, it did not provide a clear peak. The main effect of Duration was not significant (*p* = .64), but there was a significant Duration ×Time interaction with a small effect, *F*(38, 1748) = 2.31, p < 0.001, η_p_^2^ = .05. This interaction was dominated by (1) a steeper increase/decrease for Short relative to Medium for the increase from time bins 4 to 5/5 to 6, respectively, *F*(1, 46) = 6.12, *p* < 0.05, η_p_^2^ = .12; *F*(1, 46) = 4.45, *p* < 0.05, η_p_^2^ = .09, respectively, due to an earlier peak for Short relative to Medium (time point 5 *vs*. 6, respectively), and (2) a steeper increase between Medium and Long from time bins 4 to 5, *F*(1, 46) = 4.99, *p* < 0.05, η_p_^2^ = .10, due to a peak at time point 6 for medium versus 8 for long. Therefore, these effects were caused by staggered peaks/increasing extensions of activation for the Short, Medium and Long conditions, respectively, meeting experimental validity, but with a small effect size. This provides weaker evidence for a functional brain network reliably *deactivating* visual perception regions in response to auditorily presented stimuli. No main effects or interactions involving Group were significant (all *p*s > .05; see Figure 2C).

### 3.2 Melbourne and Utrecht hallucinations merged (Short/Long)

For the merged analysis of the Melbourne and Utrecht data, whereby voice hearers indicated the start and end of hallucinations by button press or ball squeeze/release, respectively, 3 components were extracted from the task-related variance in BOLD signal, as determined by examining the scree plot ^[38, 39]^. Component 1 did not show temporal or spatial validity for the Melbourne or Utrecht data, as the HDR shape was not plausible for either site, and the brain images did not show recognizable anatomical patterns, so Component 1 is not reported here (see supplementary Figure S3). Component 3 did not reveal a biologically plausible HDR shape for the Utrecht data (*p* = .05), but did for the Melbourne data, so it is reported here for the Melbourne data only, with the Utrecht data Component 3 reported in the supplementary material (Figure S4). Component 2 matched templates for the sensorimotor (response) network ^[30, Figure 7 and Table 6, 31]^, and exhibited plausible HDR shapes for both sites, meeting criteria for both spatial and temporal validity for both sites, so is reported below.

#### 3.2.1 Component 2: Sensorimotor (Response) Network

The anatomical regions associated with Component 2 are outlined in Figure 3A, and the anatomical description of Component 2 is presented in Table S4. Activation on this network involved bilateral pre- and post-central gyri (BAs 3, 4, 6), juxtapositional lobule cortex, and insular cortex (BA 47), which are all typical for the motor response network regions based on comparison with previous exemplar images _[30, Figure 7 and Table 6, 31]_.

**Figure 3:**
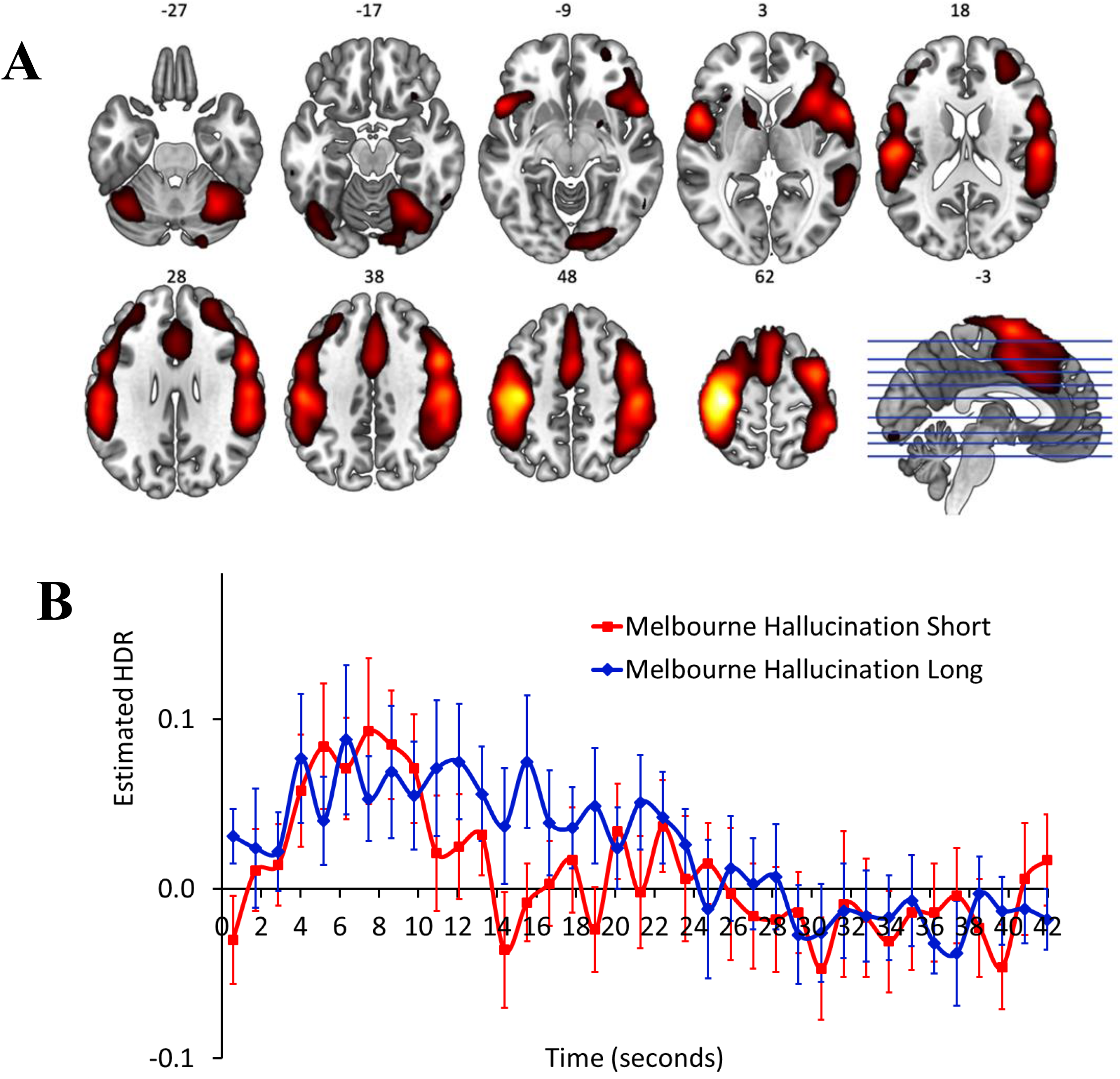
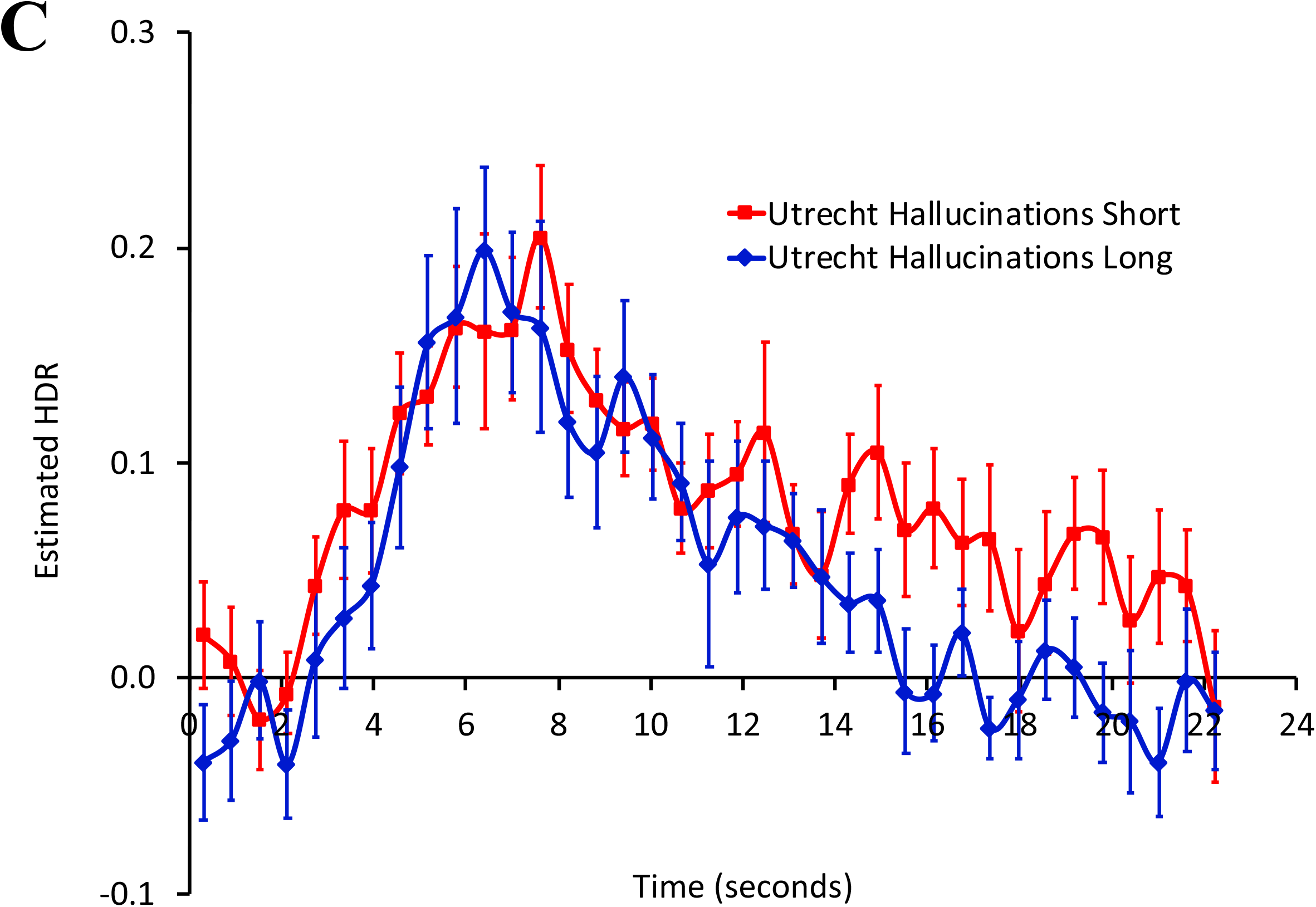
One-handed response (RESP) network, from the Melbourne and Utrecht patient hallucinations (Short/Long) merged analysis. A (top): dominant 20% of component loadings for Component 2, proposed one-handed response (RESP) network, from the Melbourne and Utrecht patient hallucinations (Short/Long) merged analysis. MNI Z-axis coordinates are displayed. Images are displayed in neurological convention (left is left). Red/yellow = positive loadings, positive threshold = 0.10, max = 0.30. B (middle): mean finite impulse response (FIR)-based predictor weights plotted as a function of post-stimulus time (TR = 610ms) and condition (averaged over participants, error bars are standard errors) for Melbourne data. C (bottom): mean finite impulse response (FIR)-based predictor weights plotted as a function of post-stimulus time (TR = 1150ms) and condition (averaged over participants, error bars are standard errors) for Utrecht data.

Figure 3B displays the estimated HDR shape for Component 2 from the Melbourne hallucinations data, for which a significant effect of Time was found, *F*(36, 396) = 2.20, *p* < 0.001, η_p_^2^ = .17. However, there was no significant effect involving duration (*p* > 0.25) suggest that this HDR shape shows temporal validity, but not experimental validity.

For the Utrecht hallucinations data (HDR shown in Figure 3C), a significant effect was found for Time *F*(36, 504) = 8.41, *p* < 0.001, η_p_^2^ = .38, showing reliability of HDR shape over participants. However, the absence of effects involving Duration (*ps* > 0.10) suggests a failure of experimental validity (see Table 4). Therefore, the pattern of activation in the response network of Utrecht participants also does not align with duration of reported hallucinations.

#### 3.2.2 Component 3: Focus on Visual Features

The anatomical regions associated with Component 3 are outlined in Figure 4A, and the anatomical description of Component 3 is presented in Table S5. The negative loadings matched the Focus on Visual Features (FVF) network, characterized by bilateral deactivation in occipital areas such as the occipital pole (xyz: 30, -91, -8; -27, -94, 1), which is known to deactivate when the visual details of the task are not relevant to response _[31, FVF, 37, Figure S2, 40, Figure 5.60]_.

**Figure 4:**
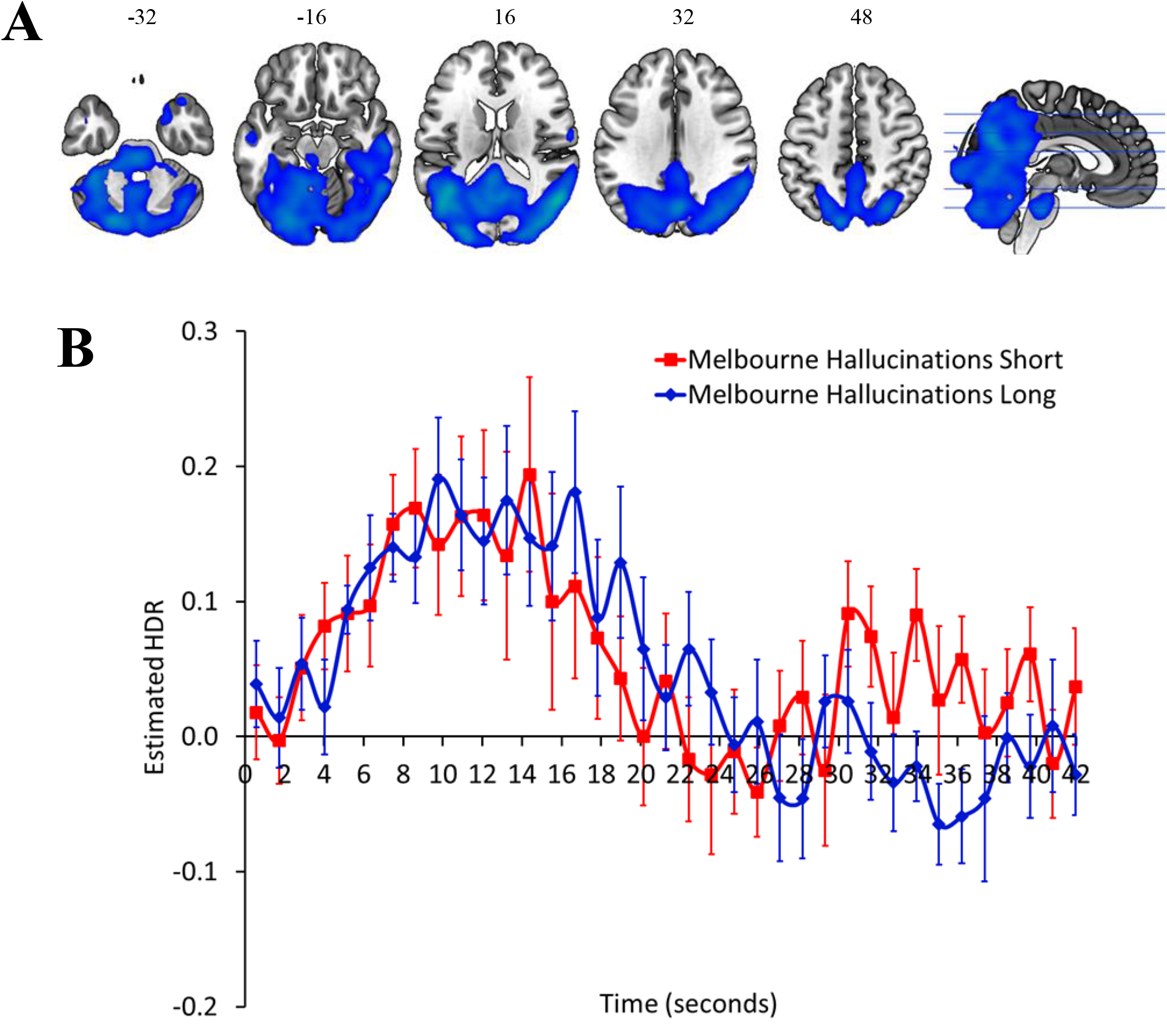
Focus on Visual Features (FVF) network, from the Melbourne (Utrecht n.s. so presented in S9) patient hallucinations (Short/Long) merged analysis. A (top): dominant 20% of component loadings for Component 3, proposed Focus on Visual Features (FVF) network, from the Melbourne (Utrecht n.s. so presented in S9) patient hallucinations (Short/Long) merged analysis. MNI Z-axis coordinates are displayed. Images are displayed in neurological convention (left is left). Blue/Green = negative loadings, negative threshold = -0.10, min = -0.18. B (bottom): mean finite impulse response (FIR)-based predictor weights plotted as a function of post-stimulus time (TR = 610ms) and condition (averaged over participants, error bars are standard errors) for Melbourne data.

Figure 4B displays the estimated HDR shape for Component 3 for the Melbourne sample. For the Melbourne data, there was a biologically plausible HDR shape, and a significant effect of Time *F*(36, 396) = 4.10, *p* < 0.05, η_p_^2^ = .27, in the absence of significant effects involving Duration (all *p*s > .1). There was not a biologically plausible HDR shape for the Utrecht data (*p* = .05; see Figure S4).

## 4. Discussion

In the current study, fMRI data were collected during AVH capture based on on-line self-reports at two sites (Melbourne and Utrecht), and merged, using fMRI-CPCA, a multidimensional analysis technique that can extract brain networks optimized to be predictable from the precise timing of the reported hallucinatory experiences. This study aimed to determine whether the functional brain networks that are detectable based on the timing of the reported experience of AVHs are underlying the hallucinatory experience itself, the experimental method of indicating the onset of the perceived hallucinatory experience (e.g., a motor response), or other cognitive events. The analyses were set up with clear criteria for spatial validity, temporal validity, and experimental validity, such that BOLD signal associated with hallucinations could be proven. From the Melbourne dataset, strong duration-dependent signal for radio speech perception was observed, which exclusively matched the template for the auditory perception network. Subsequently, in the merged analysis of the Melbourne and Utrecht data sets, a duration-dependent signal was not observed for hallucinations experiences; however, the retrieved network matched the sensorimotor (response) network. From this study, it appears that fMRI retrieved the brain network for generating responses indicating the start and end of an experienced hallucination, but failed to detect one associated with the hallucination itself.

The Table 4 Experimental Validity (Duration ×Time interaction) column shows that only radio speech elicited BOLD signal which was duration dependent, but varying durations of hallucination events were not detected by fMRI. The hallucinations blocks did reliably elicit a HDR shape though, and when this involved activation (not deactivation), this always conformed closely to the known response network (Merged: Melbourne Hallucinations C2, and Merged: Utrecht Hallucinations C2). A clear response network did not emerge for Melbourne radio speech alone, but response network regions such as left dominated precentral and postcentral gyri (xyz: -36, -19, 64; -45, -28, 49; respectively) ^[30, Figure 7 and Table 6, 31]^ were included on the Auditory Perception component.

These results are difficult to reconcile with the many previous neuroimaging studies and meta-analyses have identified brain regions showing activation during AVHs as auditory perception related ^[9, 20-22, 41]^. In Table S6 we group together the brain regions found to be involved during AVHs from three meta-analyses ^[20-22]^, in comparison to brain regions concluded to be implicated as part of the response network from the current study, and another study analyzed using fMRI-CPCA with an empirically derived response network ^[30]^. From Table S6, it can be seen that many brain regions implicated for AVHs overlap with the response network observed in component 2 from the Melbourne and Utrecht patient hallucinations (S/L) analysis. For example, the left insula (x y z peak near: -42 4 -2) has been shown to be implicated in AVH by the meta-analyses ^[20-22]^, however in the current analysis and Sanford el al. (2020), this region is considered to be a part of the response network. Similarly, activation near peaks in the right and left post central gyrus is seen in some of the meta-analyses and the response network identified form this analysis. In regards to the STG, all three meta-analyses have shown peaks of activation in this region ^[20-22]^. Although peak activation was not observed in this region in the response network from Sanford et al. (2020), peak activation was observed in adjacent brain areas to the STG, in the left central opercular cortex (x y z peak: -54 -19 16) and the right inferior frontal gyrus (x y z peak: 57 14 -2) from component 2 of the Melbourne and Utrecht patient hallucinations (S/L) analysis. However, local activation near to the STG on its own is not evidence for an underlying voice perception network in the absence for a duration-dependent signal in the HDR for hallucinations (absence of experimental validity, see Table 4 rightmost column).

The auditory perception network (Component 2, Melbourne patient/control radio speech S/M/L) is observed in many tasks as a basic sensory process activating when perceiving auditory stimuli ^[30]^. Therefore, activity in this network was expected as a result of the radio speech clips played for this analysis. The auditory perception network observed in component 2 of the Melbourne patient control radio (S/M/L) analysis achieved spatial, temporal, and experimental validity. Component 3 (Focus on Visual Features) from the Melbourne patient/control external radio (S/M/L) analysis also achieved spatial and temporal validity, although Component 2 (Auditory Perception) appeared better formed spatially and temporally.

This study was subject to some key limitations. First, the average duration of hallucinations experienced by participants varied greatly, with some participants reporting many hallucinations less than 3s long, and others having relatively long hallucinations. The method for self-reporting hallucinations may vary between individuals, with some reporting more frequently (e.g., between individual words), while others report rarely (e.g., after longer sentences), therefore making it very difficult to average over trials within participants. This inconsistency between participants may further make it more difficult to recognize functional brain networks involved in hallucinations with fMRI-CPCA, as the BOLD signal would be varying for each type of button press. In addition, the decisions regarding when to respond for the onset and offset of voices can be potentially demanding from an attention aspect.

## 5. Conclusion

In this fMRI study, the auditory perception network revealed a speech-duration-dependent HDR signal when radio clips were heard, but under no conditions were duration-dependent HDRs elicited during online-reported hallucinations. In contrast, the response network was found to underlie the button press or squeeze response when analyzing the hallucinations from merging the Utrecht and Melbourne datasets together. Therefore, no brain networks were demonstrated to be sensitive to the experience of hallucinations themselves, because duration-dependent fMRI signal was not observed in any of the components. Since responses are perfectly confounded with hallucination start and end, there is no evidence that online reporting fMRI-paradigms can detect brain networks involved in hallucinations over and above response processes. This does not imply that neurostimulation methods targeting the STG are invalid ^[42-45]^, or that hallucinations do not involve the STG, but simply suggests that either fMRI cannot detect hallucinations, or different designs may be required to indicate the onset and offset of hallucinations.

## Author Contributions

M.H., S.C., P.S., W.W., R.J., I.S., and S.L.R. collected initial data. K.G. and T.W. conceived the current study. K.G., C.P., A.C., W.S., and H.M. received and analyzed data. C.P., M.R., A.C., N.S., L.A., W.S., M.M., R.J., I.S., and S.L.R. provided intellectual contributions and/or comments on the manuscript. K.G. and T.W. wrote the first drafts of the paper.

## Supplementary Material Titles

### Methods

#### Data Analysis

***fMRI-CPCA and Repeated measures ANOVA***

***Classification of Brain Networks***

#### Timing

***Melbourne patient hallucinations (Short/Medium/Long)***

***Utrecht patient hallucinations (Short/Long)***

### Results

**Melbourne only hallucinations (Short/Medium/Long)**

**Utrecht only hallucinations (Short/Long)**

***Component 1: Response Network***

### Recommendations

**References**

**Supplementary Table S1**

*fMRI parameters*

**Supplementary Table S2**

*Cluster volumes for the most extreme 10% of Component 2 loadings (Auditory Perception Network) from the Melbourne patient control external (S/M/L) analysis, with anatomical labels, Brodmann’s areas, and MNI coordinates for the peak of each sub-cluster. Clusters smaller than 270 mm*^*3*^ *were omitted*.

**Supplementary Table S3**

*Cluster volumes for the most extreme 10% of Component 3 (Focus on Visual Features/Auditory Perception) loadings from the Melbourne patient control radio (S/M/L) analysis, with anatomical labels, Brodmann’s areas, and MNI coordinates for the peak of each sub-cluster. Clusters smaller than 270 mm*^*3*^ *were omitted*.

**Supplementary Table S4**

*Cluster volumes for the most extreme 10% of Component 2 loadings (RESP) from the Melbourne and Utrecht patient hallucinations (S/L) analysis, with anatomical labels, Brodmann’s areas, and MNI coordinates for the peak of each sub-cluster. Clusters smaller than 270 mm*^*3*^ *were omitted*.

**Supplementary Table S5**

*Cluster volumes for the most extreme 10% of Component 3 loadings (Focus on Visual Features) from the Melbourne and Utrecht patient hallucinations (S/L) analysis, with anatomical labels, Brodmann’s areas, and MNI coordinates for the peak of each sub-cluster. Clusters smaller than 270 mm*^*3*^ *were omitted*.

**Supplementary Table S6**

*Brain regions involved during motor responses and AVHs in meta analyses, with the coordinates of peak activation (in MNI atlas space) (Jardi et al*., *2011; Kompus et al*., *2011; Sanford et al*., *2020; Zmigrod et al*., *2016)*.

**Supplementary Figure S1:** Graphical Overview of fMRI-CPCA mathematical equations.

**Supplementary Figure S2:** *Melbourne patient/control radio speech (Short/Medium/Long): Component 1. A (top): dominant 20% of component loadings for Component 1, from the Melbourne patient radio speech (Short/Medium/Long) analysis. MNI Z-axis coordinates are displayed. Images are displayed in neurological convention (left is left). Red/yellow = positive loadings, positive threshold = 0*.*13, max = 0*.*21. B (middle): mean finite impulse response (FIR)-based predictor weights plotted as a function of post-stimulus time bin (TR = 1150ms) and condition (averaged over participants, error bars are standard errors)*.

**Supplementary Figure S3:** *Melbourne/Utrecht merged hallucinations (S/L) Component 1. B: Melbourne. C: Utrecht. A (top): dominant 20% of component loadings for Component 1, from the Melbourne and Utrecht patient hallucinations (Short/Long) analysis. MNI Z-axis coordinates are displayed. Images are displayed in neurological convention (left is left). Blue/green = negative loadings, negative threshold = -0*.*12, min = -0*.*18. B (middle): mean finite impulse response (FIR)-based predictor weights plotted as a function of post-stimulus time (TR = 610ms) and condition (averaged over participants, error bars are standard errors) for Melbourne data. C (bottom): mean finite impulse response (FIR)-based predictor weights plotted as a function of post-stimulus time (TR = 1150ms) and condition (averaged over participants, error bars are standard errors) for Utrecht data*.

**Supplementary Figure S4**. *Melbourne/Utrecht merged hallucinations (S/L) Component 3 (Focus on Visual Features): Utrecht. Mean finite impulse response (FIR)-based predictor weights plotted as a function of post-stimulus time (TR = 1150ms) and condition (averaged over participants, error bars are standard errors) for Utrecht data*.

**Supplementary Figure S5:** *Melbourne only hallucinations Component 1. A (top): dominant 20% of component loadings for Component 1, from the Melbourne patient hallucinations (S/M/L) analysis. MNI Z-axis coordinates are displayed. Images are displayed in neurological convention (left is left). Red/yellow = positive loadings, positive threshold = 0*.*13, max = 0*.*21. B (bottom): mean finite impulse response (FIR)-based predictor weights plotted as a function of post-stimulus time bin (TR = 1150ms) and condition (averaged over participants, error bars are standard errors)*.

**Supplementary Figure S6:** *Melbourne alone hallucinations Component 2. A (top): dominant 20% of component loadings for Component 2, proposed FVF network, from the Melbourne alone hallucinations (S/M/L) analysis. MNI Z-axis coordinates are displayed. Images are displayed in neurological convention (left is left). Blue/green = negative loadings, negative threshold = -0*.*13, min = -0*.*21. B (bottom): mean finite impulse response (FIR)-based predictor weights plotted as a function of post-stimulus time (TR = 1150ms) and condition (averaged over participants, error bars are standard errors)*.

**Supplementary Figure S7:** *Melbourne alone hallucinations Component 3. A (top): dominant 20% of component loadings for Component 3, from the Melbourne patient internal (S/M/L) analysis. MNI Z-axis coordinates are displayed. Images are displayed in neurological convention (left is left). Red/yellow = positive loadings, positive threshold = 0*.*10, max = 0*.*25. B (bottom): mean finite impulse response (FIR)-based predictor weights plotted as a function of post-stimulus time bin (TR = 1150ms) and condition (averaged over participants, error bars are standard errors)*.

**Supplementary Figure S8:** *Melbourne alone hallucinations Component 4. A (top): dominant 20% of component loadings for Component 4, from the Melbourne patient internal (S/M/L) analysis. MNI Z-axis coordinates are displayed. Images are displayed in neurological convention (left is left). Blue/green = negative loadings, negative threshold = -0*.*10, min = -0*.*24. B (bottom): mean finite impulse response (FIR)-based predictor weights plotted as a function of post-stimulus time bin (TR = 1150ms) and condition (averaged over participants, error bars are standard errors)*.

**Supplementary Figure S9:** *Utrecht alone hallucinations (S/L) Component 1. A (top): dominant 20% of component loadings for Component 1, proposed two-handed response (2RESP) network, from the Utrecht patient internal (S/L) analysis. MNI Z-axis coordinates are displayed. Images are displayed in neurological convention (left is left). Red/yellow = positive loadings, positive threshold = 0*.*10, max = 0*.*32. B (bottom): mean finite impulse response (FIR)-based predictor weights plotted as a function of post-stimulus time (TR = 610ms) and condition (averaged over participants, error bars are standard errors)*.

**Supplementary Figure S10:** *Utrecht alone hallucinations (S/L) Component 2. A (top): dominant 20% of component loadings for Component 2, from the Utrecht alone patient hallucinations (Short/Long) analysis. MNI Z-axis coordinates are displayed. Images are displayed in neurological convention (left is left). Red/yellow = positive loadings, positive threshold = 0*.*09, max 0*.*17. B (bottom): mean finite impulse response (FIR)-based predictor weights plotted as a function of post-stimulus time (TR = 610ms) and condition (averaged over participants, error bars are standard errors)*.

## References

1. Bauer SM, Schanda H, Karakula H, Olajossy-Hilkesberger L, Rudaleviciene P, Okribelashvili N, Chaudhry HR, Idemudia SE, Gscheider S, Ritter K and Stompe T. Culture and the prevalence of hallucinations in schizophrenia. Comprehensive Psychiatry 2011; 52: 319–325 [DOI: https://doi.org/10.1016/j.comppsych.2010.06.008]

2. Lim A, Hoek HW, Deen ML, Blom JD, Bruggeman R, Cahn W, de Haan L, Kahn RS, Meijer CJ, Myin-Germeys I, van Os J and Wiersma D. Prevalence and classification of hallucinations in multiple sensory modalities in schizophrenia spectrum disorders. Schizophrenia Research 2016; 176: 493–499 [DOI: https://doi.org/10.1016/j.schres.2016.06.010]

3. Rapin L, Loevenbruck H, Dohen M, Metzak PD, Whitman JC and Woodward TS. Hyperintensity of functional networks involving voice-selective cortical regions during silent thought in schizophrenia. Psychiatry Research: Neuroimaging 2012; 110–117 [DOI: doi:10.1016/j.pscychresns.2011.12.014.]

4. Ford JM, Roach BJ, Faustman WO and Mathalon DH. Synch before you speak: auditory hallucinations in schizophrenia. American Journal of Psychiatry 2007; 164: 458–466 [PMID: 17329471]

5. Northoff G and Qin P. How can the brain’s resting state activity generate hallucinations? A ‘resting state hypothesis’ of auditory verbal hallucinations. Schizophrenia Research 2011; 127: 202–14 [PMID: 21146961 DOI: S0920-9964(10)01638-5 [pii] 10.1016/j.schres.2010.11.009]

6. Lavigne K and Woodward TS. Hallucination-and speech-specific hypercoupling in frontotemporal auditory and language networks in schizophrenia using combined task-based fMRI data: an fBIRN study. Human Brain Mapping 2018; 39: 1582–1595

7. Lavigne KM, Rapin LA, Metzak PM, Whitman JC, Jung K, Dohen M, Loevenbruck H and Woodward TS. Left-dominant temporal-frontal hypercoupling in schizophrenia patients with hallucinations during speech perception. Schizophrenia Bulletin 2015; 41: 259–267

8. Suzuki M, Yuasa S, Minabe Y, Murata M and Kurachi M. Left superior temporal blood flow increases in schizophrenic and schizophreniform patients with auditory hallucination: a longitudinal case study using 123I-IMP SPECT. Eur Arch Psychiatry Clin Neurosci 1993; 242: 257–61 [PMID: 8499493]

9. Sommer IE, Diederen KM, Blom JD, Willems A, Kushan L, Slotema K, Boks MP, Daalman K, Hoek HW, Neggers SF and Kahn RS. Auditory verbal hallucinations predominantly activate the right inferior frontal area. Brain 2008; 131: 3169–77 [PMID: 18854323]

10. Hoffman RE, Hawkins KA, Gueorguieva R, Boutros NN, Rachid F, Carroll K and Krystal JH. Transcranial magnetic stimulation of left temporoparietal cortex and medication-resistant auditory hallucinations. Arch Gen Psychiatry 2003; 60: 49–56

11. Vercammen A, Knegtering H, Liemburg EJ, den Boer JA and Aleman A. Functional connectivity of the temporo-parietal region in schizophrenia: effects of rTMS treatment of auditory hallucinations. J Psychiatr Res 2010; 44: 725–31 [PMID: 20189190]

12. Jardri R, Delevoye-Turrell Y, Lucas B, Pins D, Bulot V, Delmaire C, Thomas P, Delion P and Goeb JL. Clinical practice of rTMS reveals a functional dissociation between agency and hallucinations in schizophrenia. Neuropsychologia 2009; 47: 132–8 [PMID: 18771675 DOI: S0028-3932(08)00334-5 [pii] 10.1016/j.neuropsychologia.2008.08.006]

13. Ford JM and Hoffman RE. Functional brain imaging of auditory hallucinations: from self-monitoring deficits to co-opted neural resources. The Neuroscience of Hallucinations 2013; ?

14. Hoffman R. Comment on Vercammen, Knegtering, den Boer, Liemburg, & Aleman submitted 29 April, 2010. Schizophrenia Research Forum 2010; Comment

15. David AS. The neuropsychological origin of auditory hallucinations. The neuropsychology of schizophrenia 1994; 269–313

16. Curcic-Blake B, Ford JM, Hubl D, Orlov ND, Sommer IE, Waters F, Allen P, Jardri R, Woodruff PW, David O, Mulert C, Woodward TS and Aleman A. Interaction of language, auditory and memory brain networks in auditory verbal hallucinations. Progress in neurobiology 2017; 148: 1–20 [PMID: 27890810 DOI: 10.1016/j.pneurobio.2016.11.002]

17. Allen P, Larøi F, McGuire PK and Aleman A. The hallucinating brain: a review of structural and functional neuroimaging studies of hallucinations. Neuroscience & Biobehavioral Reviews 2008; 32: 175–91 [PMID: 17884165]

18. Moritz S and Laroi F. Differences and similarities in the sensory and cognitive signatures of voice-hearing, intrusions and thoughts. Schizophr Res 2008; 102: 96–107 [PMID: 18502102]

19. Bentall RP. The illusion of reality: A review and integration of psychological research on hallucinations. Psychological Bulletin 1990; 107: 82–95

20. Kompus K, Westerhausen R and Hugdahl K. The “paradoxical” engagement of the primary auditory cortex in patients with auditory verbal hallucinations: A meta-analysis of functional neuroimaging studies. Neuropsychologia 2011; 49: 3361–3369 [DOI: http://dx.doi.org/10.1016/j.neuropsychologia.2011.08.010]

21. Zmigrod L, Garrison JR, Carr J and Simons JS. The neural mechanisms of hallucinations: A quantitative meta-analysis of neuroimaging studies. Neuroscience & Biobehavioral Reviews 2016; 69: 113–123 [DOI: https://doi.org/10.1016/j.neubiorev.2016.05.037]

22. Jardri R, Pouchet A, Pins D and Thomas P. Cortical activations during auditory verbal hallucinations in schizophrenia: a coordinate-based meta-analysis. American Journal of Psychiatry 2011; 168: 73–81 [PMID: 20952459 DOI: appi.ajp.2010.09101522 [pii] 10.1176/appi.ajp.2010.09101522]

23. Leroy A, Foucher JR, Pins D, Delmaire C, Thomas P, Roser MM, Lefebvre S, Amad A, Fovet T, Jaafari N and Jardri R. fMRI capture of auditory hallucinations: Validation of the two-steps method. Human brain mapping 2017; 38: 4966–4979 [PMID: 28660668 DOI: 10.1002/hbm.23707]

24. Fovet T, Yger P, Lopes R, Pierrefeu Ad, Duchesnay E, Houenou J, Thomas P, Szaffarczyk S, Domenech P and Jardri R. Decoding activity in Broca’s area predicts the occurrence of auditory hallucinations across subjects. bioRxiv 2021;

25. Jardri R, Thomas P, Delmaire C, Delion P and Pins D. The neurodynamic organization of modality-dependent hallucinations. Cerebral cortex (New York, N.Y. : 1991) 2013; 23: 1108–17 [PMID: 22535908 DOI: 10.1093/cercor/bhs082]

26. Beck AT and Rector NA. A Cognitive Model of Hallucinations. Cognitive Therapy and Research 2003; 27: 19–52 [DOI: 10.1023/A:1022534613005]

27. Friston K. Statistical parametric mapping. Functional neuroimaging: Technical foundations 2002; 1–74

28. Woodward TS, Cairo TA, Ruff CC, Takane Y, Hunter MA and Ngan ETC. Functional connectivity reveals load dependent neural systems underlying encoding and maintenance in verbal working memory. Neuroscience 2006; 139: 317–325

29. Metzak PD, Feredoes E, Takane Y, Wang L, Weinstein S, Cairo T, Ngan ETC and Woodward TS. Constrained principal component analysis reveals functionally connected load-dependent networks involved in multiple stages of working memory. Human Brain Mapping 2011; 32: 856–871

30. Sanford N, Whitman JC and Woodward TS. Task merging for finer separation of functional brain networks in working memory. Cortex 2020; 125: 246–271

31. Percival CM, Zahid HB and Woodward TS. Set of task-based functional brain networks derived from averaging results of multiple fMRI-CPCA studies. 2020; [DOI: https://doi.org/10.5281/zenodo.4274397]

32. Woodward TS, Zahid H and Percival CM. Brain Network Classification. 2021; UBC Invention ID: 2021–092:

33. Ribary U, Mackay AL, Rauscher A, Tipper CM, Giaschi D, Woodward TS, Sossi V, Doesburg SM, Ward LM, Herdman A, Hamarneh G, Booth BG and Moiseev A. Emerging neuroimaging technologies: Towards future personalized diagnostics, prognosis, targeted intervention and ethical challenges. Neuroethics II: Defining the Issues in Theory, Practice, and Policy 2017; 15-52

34. Fox MD, Snyder AZ, Vincent JL, Corbetta M, Van Essen DC and Raichle ME. The human brain is intrinsically organized into dynamic, anticorrelated functional networks. Proceedings of the National Academy of Sciences of the United States of America 2005; 102: 9673–8 [PMID: 15976020 DOI: 10.1073/pnas.0504136102]

35. Raichle ME, MacLeod AM, Snyder AZ, Powers WJ, Gusnard DA and Shulman GL. A default mode of brain function. Proc Natl Acad Sci U S A 2001; 98: 676–82 [PMID: 11209064]

36. Yeo BT, Krienen FM, Sepulcre J, Sabuncu MR, Lashkari D, Hollinshead M, Roffman JL, Smoller JW, Zollei L, Polimeni JR, Fischl B, Liu H and Buckner RL. The organization of the human cerebral cortex estimated by intrinsic functional connectivity. Journal of Neurophysiology 2011; 106: 1125–65 [PMID: 21653723 DOI: 10.1152/jn.00338.2011]

37. Sanford N, Whitman JC and Woodward TS. (Supplementary Data) Task merging for finer separation of functional brain networks in working memory Cortex 2020; 125: [DOI: https://doi.org/10.1016/j.cortex.2019.12.014]

38. Cattell R. The scree test for the number of factors. Multivariate Behavioural Research 1966; 1: 245–276

39. Cattell R. A Comprehensive trial of the scree and Kg criteria for determining the number of factors. Multivariate Behavioral Research 1977; 12: 289–325

40. Sanford N. Functional brain networks underlying working memory performance in schizophrenia: a multi-experiment approach. Department of Psychiatry 2019; Ph.D.:

41. Suzuki M, Yuasa S, Minabe Y, Murata M and Kurachi M. Left superior temporal blood flow increases in schizophrenic and schizophreniform patients with auditory hallucination: a longitudinal case study using 123I-IMP SPECT. European Archives of Psychiatry and Neurological Sciences 1993; 242: 257–61 [PMID: 8499493]

42. Brunelin J, Mondino M, Gassab L, Haesebaert F, Gaha L, Suaud-Chagny M-F, Saoud M, Mechri A and Poulet E. Examining Transcranial Direct-Current Stimulation (tDCS) as a Treatment for Hallucinations in Schizophrenia. American Journal of Psychiatry 2012; 169: 719–724 [PMID: 11071091 DOI: 10.1176/appi.ajp.2012.11071091]

43. Kar SK, Singh A and Prakash AJ. Neuromodulation in Schizophrenia: Relevance of Neuroimaging. Current Behavioral Neuroscience Reports 2020; 7: 139–146 [DOI: 10.1007/s40473-020-00209-2]

44. Mondino M, Jardri R, Suaud-Chagny M-F, Saoud M, Poulet E and Brunelin J. Effects of Fronto-Temporal Transcranial Direct Current Stimulation on Auditory Verbal Hallucinations and Resting-State Functional Connectivity of the Left Temporo-Parietal Junction in Patients With Schizophrenia. Schizophrenia Bulletin 2016; 42: 318–326 [DOI: 10.1093/schbul/sbv114]

45. Yang F, Fang X, Tang W, Hui L, Chen Y, Zhang C and Tian X. Effects and potential mechanisms of transcranial direct current stimulation (tDCS) on auditory hallucinations: A meta-analysis. Psychiatry Research 2019; 273: 343–349 [DOI: https://doi.org/10.1016/j.psychres.2019.01.059]

