## Supplemental Files for "Brain Networks Detectable by fMRI during On-Line Self Report of Hallucinations in Schizophrenia"

### Methods

#### Data Analysis

##### *fMRI-CPCA and Repeated measures ANOVA*

fMRI data were analyzed using constrained principal component analysis for fMRI (fMRI-CPCA) with varimax rotation [1, 2]. fMRI-CPCA is a combination of multivariate multiple regression and principal component analysis [3, 4]. Multivariate multiple regression is used to separate task-timing-predictable variance in the BOLD signal from task-unrelated variance, and principal component analysis (PCA) is applied to the task-related variance. Dominant sets of voxel-based component loadings are then interpreted spatially, alongside statistical assessment of temporal information in the estimated HDR shape. The HDR is estimated by the weights that result from regressing the component scores derived from the PCA back onto the task-timing based FIR model. The FIR model captures BOLD signal changes that are consistent over trials, and occurring approximately 20 seconds after stimulus presentation. Thus, due to the nature of the FIR model, through fMRI-CPCA, we are able to (1) identify multiple functional brain networks that are simultaneously involved in a cognitive task, (2) estimate HDR shape occurring approximately 20 seconds following stimulus presentation for each network separately, for each participant and condition, and (3) statistically test the effect of task conditions on estimated HDR shapes for each functional brain network using ANOVAs. The fMRI-CPCA application is available online, free of charge (www.nitrc.org/projects/fmricpca).

A general overview of the mathematical steps for fMRI-CPCA is provided in Figure S1, and the main three steps are described here. Step 1: multivariate least squares multiple regression of BOLD intensity values, *Z*, onto finite impulse response (FIR) model. The value 1 is placed in cells of *G* for which BOLD signal is to be estimated (based on timing information), and the value

0 in the remaining cells. The *C* matrix has condition-specific regression weights. *GC* represents the variability in *Z* predictable from *G*. Individual subject *GC* matrices are linked to create the resulting *GC* matrix. Step 2: Singular value decomposition is used to extract components in *GC* that represent temporally orthogonal functional brain networks in which coordinated BOLD activity is directly related to the hallucination timing. This also produces a *U* matrix, containing the component scores for each subject. Brain regions are observed by overlaying component loadings onto a brain template. Step 3: For each participant, component scores *(U)* are regressed onto *G* to produce predictor weights approximating the intensity of the component at each Time coded in *G*. When successfully detecting the BOLD response, predictor weights will take the shape of an HDR when plotted over time after the hallucination onset. The predictor weights can then be used for statistical analyses using repeated-measures ANOVA, since there is a separate HDR shape for each component, subject and event duration (short/medium/long). Likewise, in this current study, to test for differences between durations, subject groups, and interactions with time post stimulus, the predictor weights were submitted to an ANOVA for each of the components. Tests of sphericity were carried out, and Greenhouse-Geisser adjusted degrees of freedom were checked. Unadjusted degrees of freedom are reported, but only for effects that were also significant when adjusted degrees of freedom were used.

##### *Classification of Brain Networks*

Our in-house classification software can determine how fMRI-CPCA derived brain networks match a set of 12 candidate exemplars. It provides a *Z* statistic for a match with exemplar templates [(5)]. A MATLAB-based algorithm which outputs 12 *Z* scores for an inputted brain image. These *Z* scores indicate how closely the inputted network matches each exemplar network. This algorithm has 20-30 brain slices prototypical of each of the 12 networks digitized, and correlates the inputted network with those prototype slices, classifying based on the magnitude of the correlations, Fisher-transformed to *Z* scores. Unlike machine-learning classification algorithms, this method allows identification of hybrid networks, which can occur when networks cannot be separated. We have registered this 12-template classification algorithm with the UBC Industry Liaison Office (UILO (Invention ID 2021-092). Z values can be statistically compared to determine whether the network is a superior match to other network (Fisher Z of .8 is considered a good match, and corresponds to a Pearson's r value of .67).

**Timing**

***Melbourne patient hallucinations (Short/Medium/Long)***

For the Melbourne-only patient hallucinations analysis, timing vectors for the onset of hallucination events in the schizophrenia patients were divided into Short (1.08 - <3.69), Medium (3.69 - <5.25), and Long (5.25 - 19.00), based on indicated duration of hallucinations. Time bins for full-brain scans 1-20 following hallucination onset, as indicated by button press, were computed using a FIR basis function to monitor BOLD response (i.e., 23 seconds of post-stimulus time with TR = 1,150ms), to allow sufficient time for peak and relaxation of the HDR. After the regression and PCA steps, inspection of the scree plot of singular values indicated 4 components should be retained. Then, within subject factors of internal voice duration (*Short/Medium/Long*) and time (20 post-stimulus time bins) were examined with the resulting predictor weights, resulting in a 3 (duration) × 20 (time) ANOVA for each component. Significant effects of duration × time were further examined. Post hoc analyses of duration were carried out using polynomial contrasts, and interactions involving time were examined using repeated measures contrasts between adjacent time bins.

***Utrecht patient hallucinations (Short/Long)***

For the Utrecht-only patient hallucinations analysis, timing vectors for the onset of hallucination events in patients were divided into Short (1.02s to <7.7s) and Long (7.7s to 20.53s) applied to patients with a minimum of 4 hallucination events lasting between 1.02 and 20.6

3 seconds. The conditions for each dataset were set so that there would be nearly equal number of hallucinations in both short and long categories. Patients who did not have a minimum of 4 hallucinations events lasting between 1.02 and 20.53 seconds in duration were excluded from this analysis. Time bins for scans 1-37 following symptom onset, as indicated by squeeze response, were computed using a FIR basis function to monitor BOLD response (i.e., 22.6 seconds of post-stimulus time with TR = 610ms), to allow sufficient time for peak and relaxation of the HDR. After the regression and PCA steps, inspection of the scree plot of singular values indicated 2 components should be retained. Then, within subject factors of internal voice duration (short/long) and time (37 post-stimulus time bins) were examined with the resulting predictor weights, resulting in a 2 (duration) × 37 (time) ANOVA for the extracted component. Significant effects of duration × time were further examined. Post hoc analyses of duration were carried out using polynomial contrasts, and interactions involving time were examined using repeated measures contrasts between adjacent time bins.

#### Results

##### Melbourne only hallucinations (Short/Medium/Long)

For the Melbourne hallucinations data analyzed separately, whereby patients indicated the start and end of hallucinations by button press, 4 components were extracted from the task-related variance in BOLD signal, as determined by examining the scree plot (6, 7). However, components 1, 3, and 4 did not show temporal validity (no biologically plausible HDR shape; see

Figures S5B, S7B, and S8B, respectively), and did not show spatial validity by recognizable anatomical patterns (see Figures S5A, S7A, and S8A, respectively), so are not reported further. Component 2 showed a biologically plausible HDR shape, with a significant effect of Time, $F(19, 171) = 2.78$, $p < 0.001$, $\eta_p^2 = .24$, indicating temporal validity (see Figure S6B). It also showed a weak match to the FVF network (see Figure S6A) (5, FVF, 8, Figure S2, 9, Figure 5.60) ($Z = .54$). However, there was no effect of Duration ($p = .58$), failing experimental validity.

**Utrecht only hallucinations (Short/Long)**

For the Utrecht data analyzed separately, where patients indicated the start and end of hallucinations by ball squeeze/release, two components were extracted from the task-related variance in BOLD signal, as determined by examining the scree plot (6, 7). Component 1 exhibited a biologically plausible HDR shape, and matched the templates for the response network (1, Figure 7 and Table 6, 5, 1RESP), meeting criteria for both spatial and temporal validity, so is reported in the main manuscript text. Component 2 did not show a biologically plausible HDR shape, so is reported in Figure S10.

***Component 1: Response Network***

The anatomical regions associated with this component are outlined in Figure S9A. Activation on this network involved bilateral pre- and post-central gyri (BAs 3, 4, 6) and juxtapositional lobule cortex which are all typical motor response network regions when performing a response. Therefore, Component 1 was classified as the Response Network ($Z = .75$) based on comparison with previous exemplar images (1, Figure 7 and Table 6, 5, 1RESP). Figure S9B displays the estimated HDR shape for Component 1. The ANOVA revealed a significant effect of Time, $F(36, 504) = 4.75$, $p < 0.01$, $\eta_p^2 = .25$, suggesting temporal validity and a reliable HDR shape (see Table 4, Utrecht/Hallucinations (S/L) row). However, all effects involving Duration were not significant (all *ps* > .15), indicating that the pattern of activation in the response network did not reliably align with duration of hallucinations experienced.

#### **Recommendations**

It is possible that fMRI cannot detect hallucination, which would point to EEG or MEG as better candidates for hallucinations capture studies. However, future hallucination capture studies using fMRI would benefit from adjusted experimental designs to reveal better information about brain networks involved during the experience of hallucinations. There are a number of different ways that hallucinations capture fMRI studies would be done differently in the future to facilitate interpretation:

(1) Since it will be necessary to classify actual hallucinations into short/long, training is essential to ensure the duration of the voice matches onto the duration of the press/squeeze on/off. Clear instructions should be given to participants on how to indicate hallucination onset and offset. Participants should take part in intensive practice/training sessions involving sound files, silent thinking, and pressing for hallucinating voices to make sure it is very clear how to indicate the beginning and end of an event. It is important to avoid frequent and repeated squeezes/presses. For example, instruct the subjects to maintain the squeeze/press until there have been no voices for 2 seconds, that way, the shortest squeeze/press duration will be 2 seconds. The offset could be indicated by release of a press rather than another different press. If someone has no 2 second break from voices over the entire run, they will be squeezing the entire run, and the run will not be usable anyway, but that can't be avoided. Data should be collected on these practice/training sessions showing that participants understood the instructions.

(2) In order to avoid excessive pressing/squeezing, onset and offset of hallucinations (and radio speech or inner speech) should be indicated by onset and offset of button presses/ball squeezes, with a minimum press of 2 seconds.

(3) Alternating blocks (perhaps 2 minutes long) of pressing for hearing radio voice (e.g., 2 seconds vs. 6 seconds) inner speech (e.g., counting 2 seconds vs 6 seconds) for hallucinations (minimum 2 seconds), and pressing for counted 2 vs 6 seconds following a cue, can be compared to the same condition but without the response. 30 second rest periods can be used to allow estimation of the response network to separate from any inner speech/speech perception and hallucinations networks. There may be hallucinations when hearing radio, but the timing will match the radio not the hallucinations. Different lengths of radio sentences/inner speech/hallucinations allows networks to be elicited with staggered peaks on the HDR to ensure experimental validity.

(4) Cue the start and stop press (or squeeze), or no response, with an auditory cue, which promotes the squeeze release in the absence of internal or external speech.

(5) ITIs are very important in fMRI, and during inner speech or speech perception or button pressing, there should be a distribution of mostly short (2, 4 seconds), but a few long (6, 8 seconds) ITIs [10].

#### **References**

1. Sanford N, Whitman JC, Woodward TS (2020): Task merging for finer separation of functional brain networks in working memory. *Cortex*. 125:246-271.

2. Woodward TS, Feredoes E, Metzak PD, Takane Y, Manoach DS (2013): Epoch-specific functional networks involved in working memory. *Neuroimage*. 65:529-539.

3. Takane Y, Hunter MA (2001): Constrained principal component analysis: A comprehensive theory. *Applicable Algebra in Engineering, Communication and Computing*. 12:391-419.

4. Takane Y, Shibayama T (1991): Principal component analysis with external information on both subjects and variables. *Psychometrika*. 56:97-120.

5. Percival CM, Zahid HB, Woodward TS (2020): Set of task-based functional brain networks derived from averaging results of multiple fMRI-CPCA studies. CNoS-Lab/Woodward_Atlas. Zenodo.

6. Cattell R (1966): The scree test for the number of factors. *Multivariate Behavioural Research*. 1:245-276.

7. Cattell R (1977): A Comprehensive trial of the scree and Kg criteria for determining the number of factors. *Multivariate Behavioral Research*. 12:289-325.

8. Sanford N, Whitman JC, Woodward TS (2020): (Supplementary Data) Task merging for finer separation of functional brain networks in working memory *Cortex*. 125.

9. Sanford N (2019): Functional brain networks underlying working memory performance in schizophrenia: a multi-experiment approach. Vancouver, Canada: University of British Columbia.

10. Zarahn E, Aguirre G, D'Esposito M (1997): A trial-based experimental design for fMRI. *Neuroimage*. 6:122-138.

**Supplementary Table S1**

*fMRI parameters*

| | Scanner Type | Sequence | TR (s) | # Runs | Length of Runs (TRs) | Minutes scanned |
|---|---|---|---|---|---|---|
| Melbourne | 3T Siemens | EPI | 1.15 | 1 | 1154-1300 | 22-25 |
| Utrecht | 3T Philips | PRESTO | 0.61 | 1-3 | 800 | 8-24 |

**Supplementary Table S2**

*Cluster volumes for the most extreme 10% of Component 2 loadings (Auditory Perception Network) from the Melbourne patient control external (S/M/L) analysis, with anatomical labels, Brodmann's areas, and MNI coordinates for the peak of each sub-cluster. Clusters smaller than 270 mm³ were omitted.*

| Anatomical Label | Cluster Volume (mm²) | Brodmann's Area for Peak Locations | MNI Coordinate for Peak Locations x | y | z |
|---|---|---|---|---|---|
| | | *Positive Loadings* | | | |
| *Cluster 1: Right hemisphere* | 82701 | | | | |
| *Superior Temporal Gyrus, posterior division* | | 22 | 60 | -16 | -2 |
| *Inferior Frontal Gyrus, pars opercularis* | | 48 | 42 | 11 | 25 |
| *Precentral Gyrus* | | 6 | 51 | 2 | 43 |
| *Frontal Operculum Cortex* | | 47 | 36 | 23 | 4 |
| *Cluster 2: Left hemisphere* | 66393 | | | | |
| *Planum Temporale* | | 22 | -57 | -19 | 1 |
| *Left Cerebral White Matter* | | n/a | -24 | -7 | -8 |
| *Inferior Frontal Gyrus, pars opercularis* | | 44 | -45 | 8 | 19 |
| *Precentral Gyrus* | | 6 | -54 | 5 | 19 |
| *Cluster 3: Left hemisphere* | 12069 | | | | |
| *Precentral Gyrus* | | 4 | -36 | -19 | 64 |
| *Postcentral Gyrus* | | 3 | -45 | -28 | 49 |
| *Cluster 4: Bilateral* | 7884 | | | | |
| *Left Thalamus* | | 27 | -12 | -28 | -2 |
| *Left Thalamus* | | n/a | -12 | -19 | 4 |
| *Right Thalamus* | | n/a | 9 | -16 | 7 |
| *Cluster 5: Left hemisphere* | 3861 | | | | |
| *Juxtapositional Lobule Cortex* | | n/a | -3 | -1 | 49 |

**Supplementary Table S3**

*Cluster volumes for the most extreme 10% of Component 3 (Focus on Visual Features/Auditory Perception) loadings from the Melbourne patient control radio (S/M/L) analysis, with anatomical labels, Brodmann's areas, and MNI coordinates for the peak of each sub-cluster. Clusters smaller than 270 mm³ were omitted.*

| Anatomical Label | Cluster Volume ($mm^2$) | Brodmann's Area for Peak Locations | MNI Coordinate for Peak Locations x | y | z |
|---|---|---|---|---|---|
| | *Negative Loadings* | | | | |
| *Cluster 1: Bilateral* | 173016 | | | | |
| *Occipital Pole* | | 18 | -27 | -91 | 16 |
| *Lateral Occipital Cortex, superior division* | | 19 | -24 | -79 | 31 |
| *Precuneous Cortex* | | n/a | -6 | -70 | 22 |
| *Lateral Occipital Cortex, superior division* | | 19 | -24 | -79 | 37 |
| *Precuneous Cortex* | | 18 | -9 | -67 | 19 |
| *Intracalcarine Cortex* | | 17 | -6 | -64 | 13 |
| *Lingual Gyrus* | | 17 | -6 | -64 | 7 |
| *Lateral Occipital Cortex, superior division* | | 19 | 30 | -82 | 31 |
| *Precuneous Cortex* | | 17 | 9 | -58 | 7 |
| *Cuneal Cortex* | | 18 | 3 | -76 | 34 |
| *Cuneal Cortex* | | 19 | -9 | -88 | 31 |
| *Cuneal Cortex* | | n/a | 12 | -70 | 28 |
| *Lateral Occipital Cortex, superior division* | | 19 | 30 | -88 | 19 |
| *Lateral Occipital Cortex, superior division* | | 19 | 15 | -79 | 43 |
| *Cuneal Cortex* | | 18 | 3 | -85 | 25 |
| *Occipital Pole* | | 18 | 6 | -91 | 13 |
| *Lateral Occipital Cortex, inferior division* | | 19 | -45 | -76 | -2 |
| *Occipital Fusiform Gyrus* | | 19 | 24 | -67 | -14 |
| *Occipital Pole* | | 18 | 15 | -88 | 28 |
| *Occipital Pole* | | 17 | -9 | -100 | 10 |
| *Occipital Fusiform Gyrus* | | 19 | -27 | -73 | -17 |
| *Lateral Occipital Cortex, inferior division* | | 19 | 42 | -76 | 10 |
| *Lateral Occipital Cortex, inferior division* | | 19 | 42 | -82 | -5 |
| *Occipital Pole* | | 18 | 33 | -91 | 1 |
| *Occipital Pole* | | 18 | 24 | -97 | -2 |
| *Precuneous Cortex* | | n/a | -6 | -61 | 46 |
| *Occipital Fusiform Gyrus* | | 18 | 21 | -82 | -17 |
| *Lateral Occipital Cortex, inferior division* | | 19 | 39 | -85 | -11 |
| *Occipital Pole* | | 18 | 30 | -91 | -8 |
| *Occipital Fusiform Gyrus* | | 18 | -18 | -85 | -14 |
| *Lingual Gyrus* | | 17 | -9 | -79 | -11 |

**Supplementary Table S4**

*Cluster volumes for the most extreme 10% of Component 2 loadings (RESP) from the Melbourne and Utrecht patient hallucinations (S/L) analysis, with anatomical labels, Brodmann's areas, and MNI coordinates for the peak of each sub-cluster. Clusters smaller than 270 mm$^3$ were omitted.*

| Anatomical Label | Cluster Volume (mm$^2$) | Brodmann's Area for Peak Locations | MNI Coordinate for Peak Locations | | |
|---|---|---|---|---|---|
| | | | x | y | z |
| *Positive Loadings* | | | | | |
| *Cluster 1: Bilateral* | 201447 | | | | |
| *Precentral Gyrus* | | 4 | -36 | -22 | 61 |
| *Precentral Gyrus* | | 44 | 51 | 8 | 37 |
| *Juxtapositional Lobule Cortex* | | n/a | 0 | 2 | 70 |
| *Postcentral Gyrus* | | 3 | 51 | -22 | 43 |
| *Superior Parietal Lobule* | | 2 | 45 | -40 | 58 |
| *Middle Frontal Gyrus* | | 6 | 39 | -1 | 58 |
| *Inferior Frontal Gyrus, pars opercularis* | | 38 | 57 | 14 | -2 |
| *Precentral Gyrus* | | 6 | 36 | -4 | 61 |
| *Central Opercular Cortex* | | 48 | -54 | -19 | 16 |
| *Precentral Gyrus* | | 48 | -54 | 8 | 1 |
| *Central Opercular Cortex* | | 42 | 60 | -19 | 16 |
| *Parietal Operculum Cortex* | | 2 | 57 | -34 | 34 |
| *Superior Frontal Gyrus* | | 6 | 21 | -1 | 70 |
| *Supramarginal Gyrus, anterior division* | | 48 | -60 | -31 | 28 |
| *Superior Frontal Gyrus* | | 6 | -12 | -1 | 70 |
| *Insular Cortex* | | 48 | 36 | 20 | 4 |
| *Juxtapositional Lobule Cortex* | | n/a | 0 | 2 | 52 |
| *Precentral Gyrus* | | 6 | -54 | 5 | 37 |
| *Precentral Gyrus* | | 6 | -51 | 2 | 43 |
| *Precentral Gyrus* | | 44 | -54 | 8 | 31 |
| *Paracingulate Gyrus* | | 24 | 3 | 23 | 37 |
| *Middle Temporal Gyrus, temporooccipital part* | | 22 | 60 | -43 | 10 |
| *Cluster 2: Right hemisphere* | 7857 | | | | |
| *Occipital Fusiform Gyrus* | | 19 | 36 | -79 | -14 |
| *Cluster 3: Left hemisphere* | 2133 | | | | |
| *Temporal Occipital Fusiform Cortex* | | 37 | -39 | -61 | -23 |
| *Occipital Fusiform Gyrus* | | 19 | -39 | -76 | -20 |
| *Cluster 4: Right hemisphere* | 1809 | | | | |
| *Frontal Pole* | | 46 | 30 | 47 | 25 |
| *Middle Frontal Gyrus* | | 46 | 39 | 35 | 31 |

**Supplementary Table S5**

*Cluster volumes for the most extreme 10% of Component 3 loadings (Focus on Visual Features) from the Melbourne and Utrecht patient hallucinations (S/L) analysis, with anatomical labels, Brodmann's areas, and MNI coordinates for the peak of each sub-cluster. Clusters smaller than 270 mm³ were omitted.*

| Anatomical Label | Cluster Volume ($mm^2$) | Brodmann's Area for Peak Locations | MNI Coordinate for Peak Locations x | y | z |
|---|---|---|---|---|---|
| | | *Negative Loadings* | | | |
| *Cluster 1: Bilateral* | 210087 | | | | |
| *Occipital Pole* | | 18 | 30 | -91 | -8 |
| *Occipital Pole* | | 18 | -27 | -94 | 1 |
| *Occipital Pole* | | 17 | 21 | -100 | -2 |
| *Occipital Pole* | | 18 | -27 | -91 | 13 |
| *Lateral Occipital Cortex, superior division* | | 37 | 54 | -64 | 16 |
| *Lateral Occipital Cortex, inferior division* | | 19 | 48 | -76 | 7 |
| *Lateral Occipital Cortex, inferior division* | | 19 | 51 | -73 | 4 |
| *Lateral Occipital Cortex, inferior division* | | 19 | 42 | -79 | -11 |
| *Angular Gyrus* | | 39 | -39 | -55 | 19 |
| *Lateral Occipital Cortex, inferior division* | | 37 | 57 | -61 | 7 |
| *Lateral Occipital Cortex, inferior division* | | 37 | 54 | -67 | 4 |
| *Occipital Pole* | | 18 | -9 | -97 | -8 |
| *Occipital Fusiform Gyrus* | | 18 | -21 | -67 | -11 |
| *Lateral Occipital Cortex, superior division* | | 39 | -45 | -67 | 16 |
| *Lateral Occipital Cortex, superior division* | | 39 | -36 | -70 | 19 |
| *Temporal Fusiform Cortex, posterior division* | | 37 | -36 | -37 | -23 |
| *Lateral Occipital Cortex, superior division* | | 19 | -12 | -82 | 46 |
| *Lateral Occipital Cortex, inferior division* | | 19 | -42 | -76 | -5 |
| *Lateral Occipital Cortex, superior division* | | 19 | 45 | -76 | 16 |
| *Temporal Fusiform Cortex, posterior division* | | 37 | -39 | -40 | -26 |
| *Precuneous Cortex* | | n/a | 0 | -70 | 28 |
| *Lateral Occipital Cortex, superior division* | | 19 | 36 | -85 | 19 |
| *Occipital Fusiform Gyrus* | | 18 | -24 | -88 | -14 |
| *Occipital Fusiform Gyrus* | | 19 | -30 | -85 | -14 |
| *Angular Gyrus* | | 22 | 57 | -55 | 25 |
| *Angular Gyrus* | | 22 | 60 | -49 | 25 |
| *Intracalcarine Cortex* | | 19 | -21 | -61 | 7 |
| *Intracalcarine Cortex* | | 17 | -18 | -64 | 10 |
| *Lingual Gyrus* | | 18 | -12 | -61 | -8 |
| *Precuneous Cortex* | | 23 | 6 | -61 | 19 |
| *Occipital Pole* | | 18 | 24 | -94 | 19 |
| *Lingual Gyrus* | | 18 | -3 | -70 | -2 |

| | | | | | |
|---|---|---|---|---|---|
| *Inferior Temporal Gyrus, temporooccipital part* | | 37 | 54 | -58 | -14 |
| *Lateral Occipital Cortex, superior division* | | 39 | 42 | -76 | 28 |
| *Lateral Occipital Cortex, superior division* | | 19 | 30 | -79 | 37 |
| *Cingulate Gyrus, posterior division* | | 30 | -3 | -46 | 19 |
| *Brain-Stem* | | n/a | 0 | -28 | -32 |
| *Lateral Occipital Cortex, superior division* | | 7 | 30 | -70 | 46 |
| *Precuneous Cortex* | | 17 | -6 | -58 | 16 |
| *Precuneous Cortex* | | 23 | -15 | -67 | 22 |
| *Lateral Occipital Cortex, inferior division* | | 19 | -39 | -70 | 1 |
| *Temporal Fusiform Cortex, posterior division* | | 20 | -36 | -28 | -20 |
| *Precuneous Cortex* | | 7 | 0 | -67 | 52 |
| *No Match* | | 37 | -18 | -43 | -23 |
| *Temporal Occipital Fusiform Cortex* | | 37 | 48 | -49 | -26 |
| *Lingual Gyrus* | | 19 | 27 | -55 | -5 |
| *Precuneous Cortex* | | n/a | 3 | -49 | 40 |
| *Intracalcarine Cortex* | | n/a | 21 | -64 | 7 |
| *Lingual Gyrus* | | 18 | 12 | -64 | -5 |
| *Middle Temporal Gyrus, temporooccipital part* | | 37 | -54 | -61 | -2 |
| *Cluster 2: Right hemisphere* | 1539 | | | | |
| *Temporal Fusiform Cortex, posterior division* | | 20 | 39 | -31 | -20 |
| *Inferior Temporal Gyrus, posterior division* | | 20 | 45 | -37 | -17 |
| *Cluster 3: Right hemisphere* | 918 | | | | |
| *Middle Temporal Gyrus, posterior division* | | 20 | 57 | -19 | -17 |
| *Cluster 4: Left hemisphere* | 729 | | | | |
| *Middle Temporal Gyrus, posterior division* | | 22 | -60 | -16 | -8 |

**Supplementary Table S6**

*Brain regions involved during motor responses and AVHs in meta analyses, with the coordinates of peak activation (in MNI atlas space) (Jardri et al., 2011; Kompus et al., 2011; Sanford et al., 2020; Zmigrod et al., 2016).*

| | MNI coord (mm) | | | | MNI coord (mm) | | | | MNI coord (mm) | | | | MNI coord (mm) | | | | MNI coord (mm) | | |
|---|---|---|---|---|---|---|---|---|---|---|---|---|---|---|---|---|---|---|---|
| Melbourne_Utrecht pat hallucinations (S/L); C2 Response network | x | y | z | Sanford et al. (2020) - C1 Response network | x | y | z | Jardri et al. (2011) - AVH | x | y | z | Kompus et al. (2011) - AVH | x | y | z | Zmigrod et al. (2016) - AVH | x | y | z |
| Precentral Gyrus | 51 | 8 | 37 | Precentral gyrus | 57 | 8 | 17 | | | | | Inferior frontal gyrus | 42 | 14 | 13 | Inferior frontal gyrus | 60 | 8 | 12 |
| | | | | Frontal operculum cortex | 48 | 11 | 5 | | | | | | | | | Inferior frontal gyrus | 56 | 16 | 8 |
| | | | | | | | | | | | | | | | | Inferior frontal gyrus | 50 | 24 | 0 |
| | | | | | | | | | | | | | | | | Insula | 40 | 0 | 12 |
| Middle Frontal Gyrus | 39 | 35 | 31 | Middle fronal gyrus | 39 | 35 | 29 | | | | | Superior frontal gyrus | 27 | 42 | 28 | | | | |
| Frontal Pole | 30 | 47 | 25 | Frontal pole | 36 | 44 | 26 | | | | | | | | | | | | |
| Insular Cortex | 36 | 20 | 4 | Insula | 33 | 23 | 2 | | | | | | | | | | | | |
| Paracingulate gyrus | 3 | 23 | 7 | Paracingulate gyrus | 3 | 17 | 47 | | | | | | | | | Medial frontal gyrus | 6 | 6 | 60 |
| | | | | | | | | | | | | | | | | Medial frontal gyrus | 0 | 12 | 46 |
| Supplementary Motor Cortex | 0 | 2 | 52 | Supplementary motor area | -3 | -4 | 53 | | | | | | | | | | | | |
| Supplementary Motor Cortex | 0 | 2 | 70 | | | | | | | | | | | | | | | | |
| Postcentral Gyrus | 51 | -22 | 43 | Postcentral gyrus | 57 | -16 | 29 | | | | | Middle temporal gyrus | 59 | -45 | 11 | Postcentral Gyrus | 64 | -16 | 36 |
| Central Opercular Cortex | 60 | -19 | 16 | Postcentral gyrus | 54 | -19 | 38 | | | | | Inferior parietal lobule | 31 | -44 | 53 | Postcentral Gyrus | 60 | -22 | 44 |
| Parietal Operculum Cortex | 57 | -34 | 34 | Posterior supramarginal gyrus | 48 | -37 | 50 | | | | | | | | | | | | |
| Superior Parietal Lobule | 45 | -40 | 58 | Superior parietal lobule | 39 | -46 | 50 | | | | | | | | | | | | |
| Middle Temporal Gyrus, temporooccipital part | 60 | -43 | 10 | Posterior supramarginal gyrus | 36 | -46 | 41 | | | | | | | | | | | | |
| Middle Frontal Gyrus | 39 | -1 | 58 | Middle fronal gyrus | 33 | -1 | 59 | | | | | | | | | | | | |
| Precentral Gyrus | 36 | -4 | 61 | | | | | | | | | | | | | | | | |

| | | | | | | | | | | | | | | | | | | | |
|---|---|---|---|---|---|---|---|---|---|---|---|---|---|---|---|---|---|---|---|
| Superior Frontal Gyrus | 21 | -1 | 70 | | | | | | | | | | | | | | | | |
| | | | | | | | | | | | | | | | | Claustrum | 40 | -4 | 4 |
| | | | | Anterior internal capsule | 15 | -1 | 14 | | | | | | | | | Thalamus | 18 | -10 | 2 |
| | | | | Thalamus | 12 | -16 | 8 | | | | | | | | | | | | |
| | | | | Precuneus cortex | 9 | -67 | 50 | | | | | | | | | | | | |
| Inferior Frontal Gyrus, pars opercularis | 57 | 14 | -2 | | | | | Anterior insula | 47 | 10 | -10 | | | | | Superior Temporal Gyrus | 54 | 12 | -6 |
| | | | | | | | | Frontal operculum | 44 | 17 | -17 | | | | | Insula | 48 | 8 | -12 |
| | | | | | | | | | | | | Middle temporal gyrus | 57 | -31 | -11 | Middle Temporal Gyrus | 60 | -32 | -6 |
| No Match | 30 | -61 | -23 | Cerebellum V | 18 | -52 | -22 | | | | | Cerebellum lobule V | 21 | -45 | -24 | Cerebellum | 26 | -54 | -20 |
| Occipital Fusiform Gyrus | 36 | -79 | -14 | Cerebellum VIIIa | 18 | -64 | -49 | | | | | | | | | Cerebellum | 26 | -62 | -46 |
| | | | | Cerebellum V | 12 | -58 | -28 | | | | | | | | | Cerebellum | 16 | -56 | -20 |
| | | | | Cerebellum crus II | 6 | -73 | -34 | | | | | | | | | Cerebellum | 16 | -58 | -52 |
| | | | | Cerebellum VIIb | 3 | -67 | -49 | | | | | | | | | | | | |
| | | | | Cerebellum VI | -6 | -73 | -13 | | | | | | | | | | | | |
| | | | | Cerebellum VI | -9 | -70 | -19 | | | | | | | | | | | | |
| | | | | Lingual gyrus | -3 | -79 | -1 | | | | | | | | | | | | |
| | | | | | | | | | | | | | | | | Medial frontal gyrus | -2 | 8 | 60 |
| | | | | Insula | -30 | 20 | 5 | | | | | | | | | | | | |
| | | | | Frontal pole | -33 | 44 | 20 | | | | | | | | | | | | |
| | | | | Middle fronal gyrus | -36 | 35 | 26 | | | | | | | | | | | | |
| Precentral Gyrus | -54 | 5 | 37 | Precentral gyrus | -51 | 5 | 11 | Anerior Insula | -44 | 2 | 3 | Insula | -46 | 0 | 3 | Insula | -42 | 4 | -2 |
| Precentral Gyrus | -51 | 2 | 43 | Precentral gyrus | -54 | 8 | 29 | Precentral gyrus | -57 | 2 | 13 | | | | | Insula | -44 | 0 | 4 |
| Precentral Gyrus | -54 | 8 | 31 | Insula | -39 | -1 | 11 | Broca's convolution | -50 | 13 | 5 | | | | | Precentral gyrus | -56 | 4 | 12 |
| Precentral Gyrus | -54 | 8 | 1 | | | | | | | | | | | | | | | | |
| Superior Frontal Gyrus | -12 | -1 | 70 | | | | | | | | | | | | | | | | |
| Precentral Gyrus | -36 | -22 | 61 | Precentral gyrus | -39 | -19 | 59 | | | | | Postcentral gyrus | -51 | -26 | 43 | Postcentral Gyrus | -40 | -16 | 54 |
| | | | | Postcentral gyrus | -51 | -25 | 47 | | | | | | | | | Postcentral Gyrus | -46 | -18 | 44 |

| | | | | | | | | | | | | | | | | | | | |
|---|---|---|---|---|---|---|---|---|---|---|---|---|---|---|---|---|---|---|---|
| | | | | Superior parietal lobule | -33 | -52 | 41 | | | | | | | | | Postcentral Gyrus | -52 | -22 | 50 |
| Supramarginal Gyrus, anterior division | -60 | -31 | 28 | Central opercular cortex | -54 | -19 | 20 | | | | | | | | | | | | |
| | | | | parietal operculum cortex | -57 | -40 | 23 | | | | | | | | | | | | |
| | | | | Superior lateral occipital cortex | -12 | -70 | 50 | | | | | | | | | | | | |
| Central Opercular Cortex | -54 | -19 | 16 | | | | | Supramarginal gyrus | -55 | -20 | 14 | Superior temporal gyrus | -55 | -22 | 15 | Superior Temporal Gyrus | -58 | -46 | 20 |
| | | | | | | | | Superior and middle temporal gyri | -56 | -45 | 14 | | | | | Superior Temporal Gyrus | -60 | -56 | 20 |
| | | | | | | | | | | | | | | | | Insula | -48 | -40 | 24 |
| | | | | Thalamus | -12 | -19 | 8 | | | | | | | | | Thalamus | -12 | -20 | 4 |
| | | | | Puamen | -30 | -16 | 2 | | | | | | | | | Thalamus | -28 | -32 | 8 |
| | | | | Inferior lateral occipital cortex | -48 | -64 | 5 | | | | | | | | | | | | |
| | | | | Middle temporal gyrus, temporoccipital part | -48 | -52 | 8 | | | | | | | | | | | | |
| | | | | | | | | Hippocampus/parahippocampal gyrus | -26 | -31 | -10 | Hippocampus | -26 | -31 | 10 | | | | |
| | | | | Temporal pole | -51 | 11 | -4 | | | | | | | | | | | | |
| Temporal Occipital Fusiform Cortex | -39 | -61 | -23 | Cerebellum VI | -30 | -55 | -31 | | | | | | | | | | | | |
| Occipital Fusiform Gyrus | -39 | -76 | -20 | Cerebellum VIIb | -33 | -58 | -49 | | | | | | | | | | | | |
| | | | | | | | | | | | | | | | | Midbrain | -16 | -24 | -4 |
| | | | | | | | | | | | | | | | | Parahippocampal Gyrus | -26 | -32 | -4 |

**Supplementary Figure S1:** *Graphical Overview of fMRI-CPCA mathematical equations.*

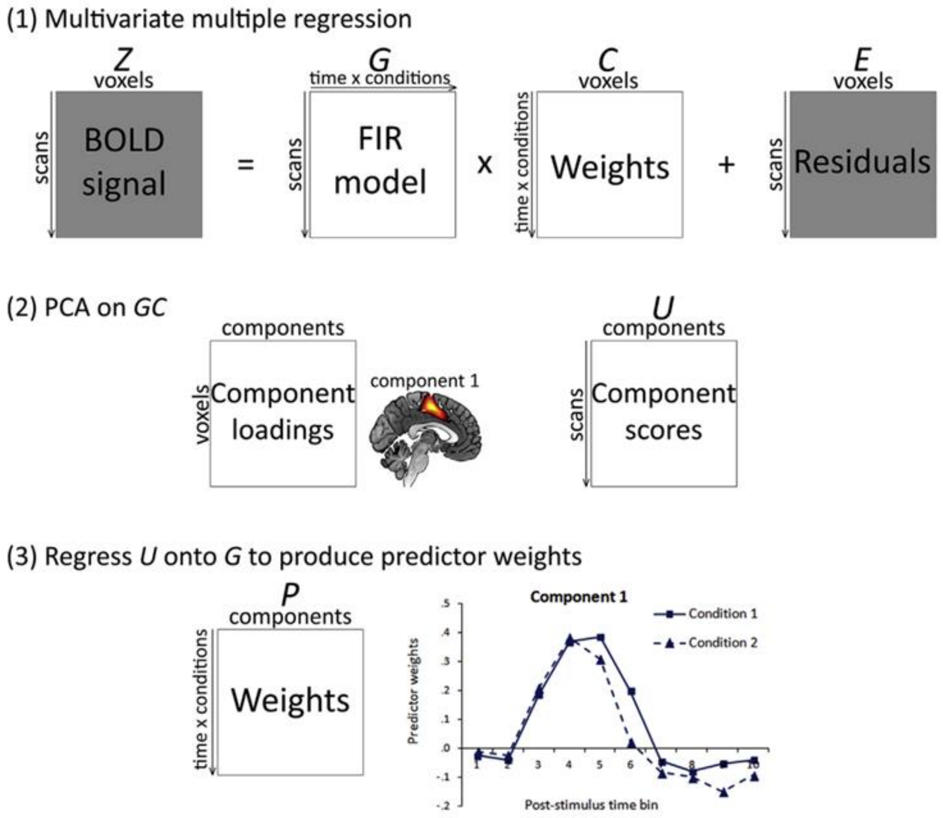

**Supplementary Figure S2:** *Melbourne patient/control radio speech (Short/Medium/Long): Component 1. A (top): dominant 20% of component loadings for Component 1, from the Melbourne patient radio speech (Short/Medium/Long) analysis. MNI Z-axis coordinates are displayed. Images are displayed in neurological convention (left is left). Red/yellow = positive loadings, positive threshold = 0.13, max = 0.21. B (middle): mean finite impulse response (FIR)-based predictor weights plotted as a function of post-stimulus time bin (TR = 1150ms) and condition (averaged over participants, error bars are standard errors).*

A

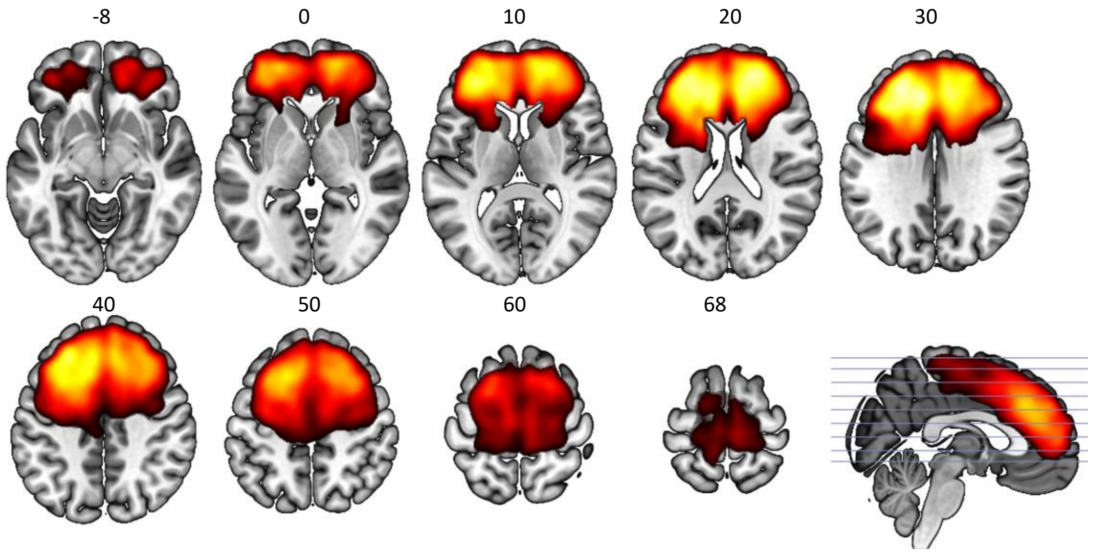


B

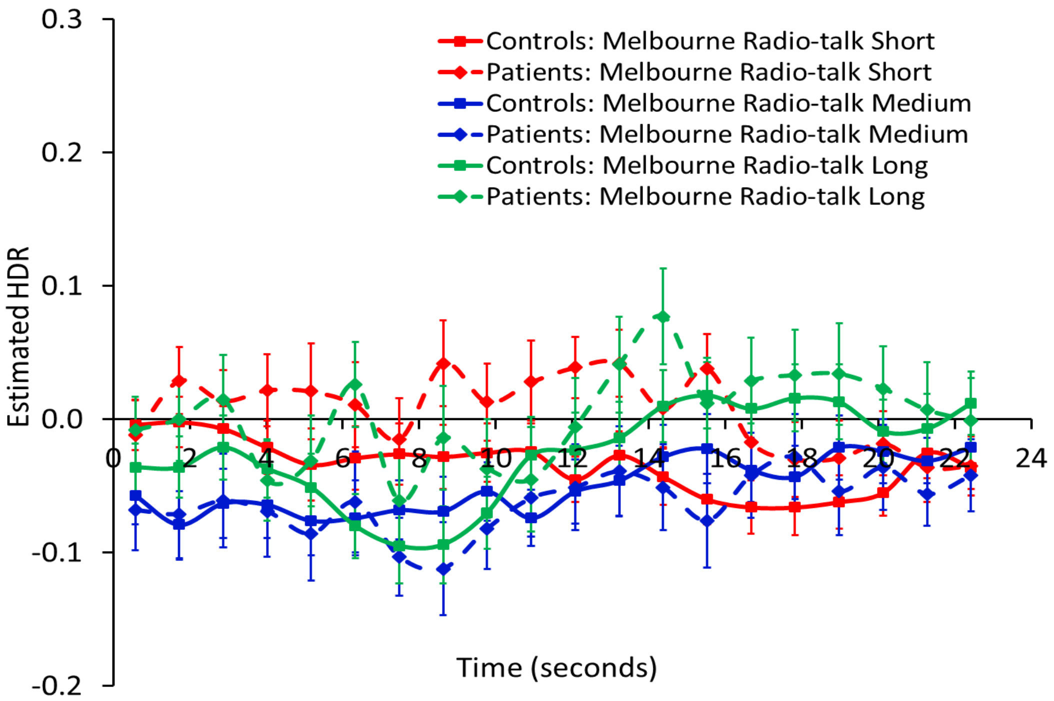

**Supplementary Figure S3:** *Melbourne/Utrecht merged hallucinations (S/L) Component 1. B: Melbourne. C: Utrecht. A (top): dominant 20% of component loadings for Component 1, from the Melbourne and Utrecht patient hallucinations (Short/Long) analysis. MNI Z-axis coordinates are displayed. Images are displayed in neurological convention (left is left). Blue/green = negative loadings, negative threshold = -0.12, min = -0.18. B (middle): mean finite impulse response (FIR)-based predictor weights plotted as a function of post-stimulus time (TR = 610ms) and condition (averaged over participants, error bars are standard errors) for Melbourne data. C (bottom): mean finite impulse response (FIR)-based predictor weights plotted as a function of post-stimulus time (TR = 1150ms) and condition (averaged over participants, error bars are standard errors) for Utrecht data.*

A

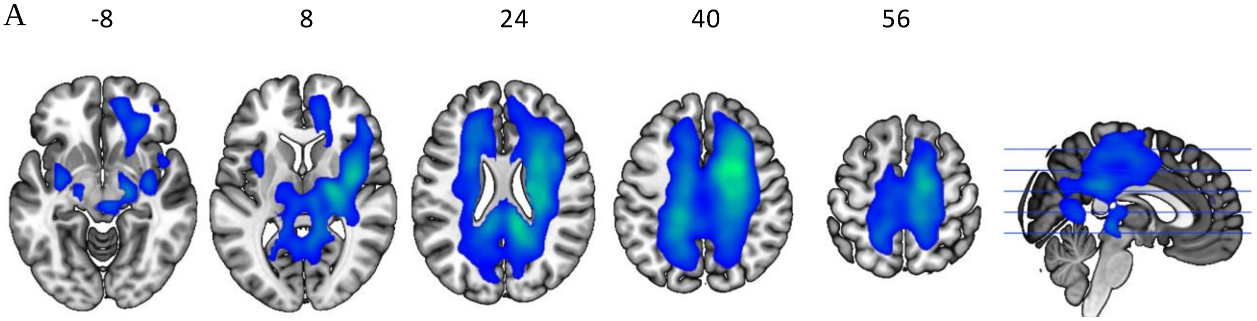


B

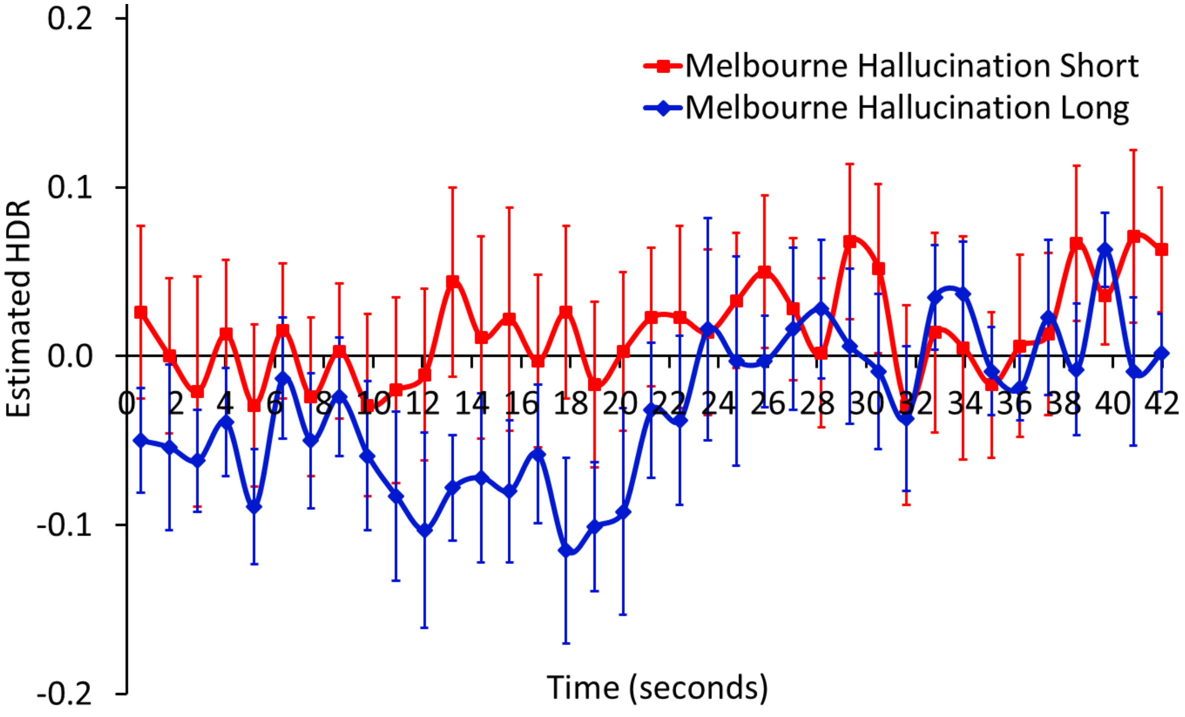

C

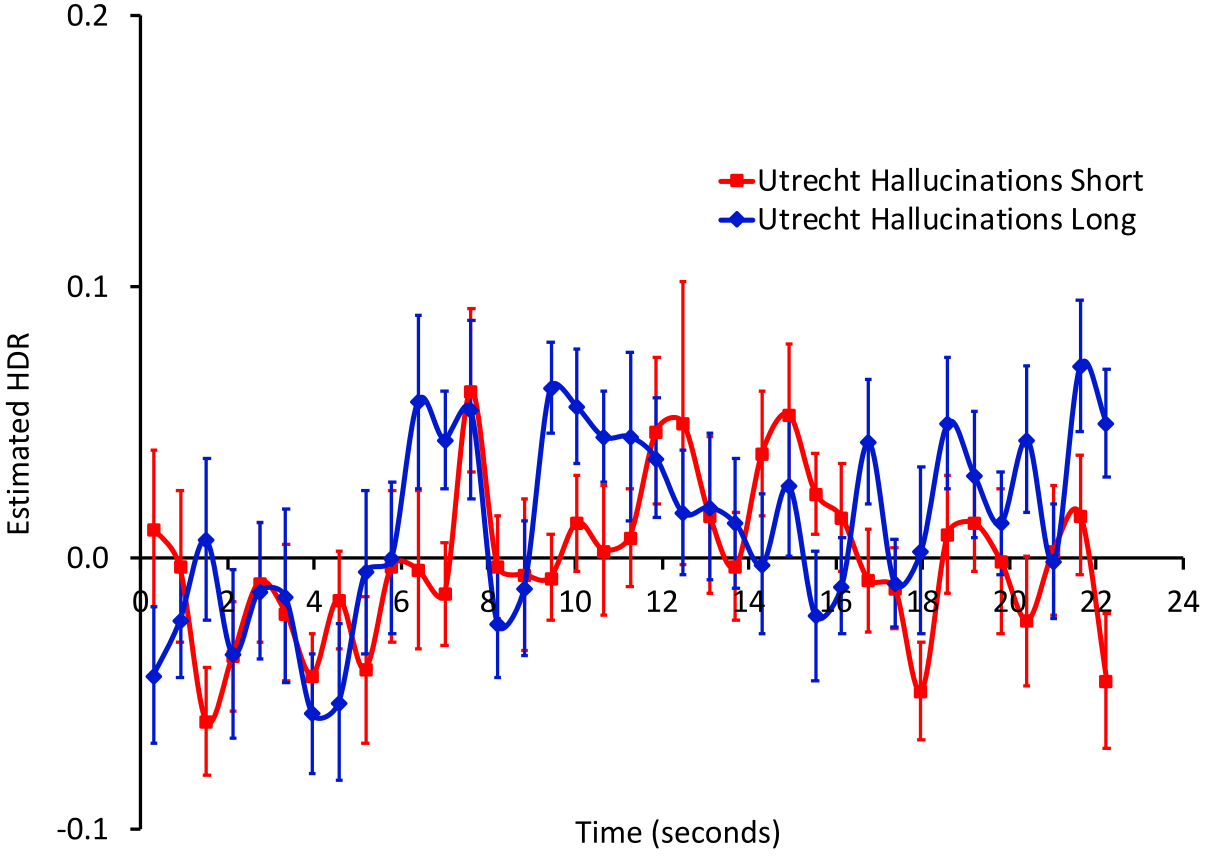
0.2
0.1
0.0
-0.1
Estimated HDR
0
2
4
6
8
10
12
14
16
18
20
22
24
Time (seconds)
Utrecht Hallucinations Short
Utrecht Hallucinations Long

**Supplementary Figure S4.** *Melbourne/Utrecht merged hallucinations (S/L) Component 3 (Focus on Visual Features): Utrecht. Mean finite impulse response (FIR)-based predictor weights plotted as a function of post-stimulus time (TR = 1150ms) and condition (averaged over participants, error bars are standard errors) for Utrecht data.*

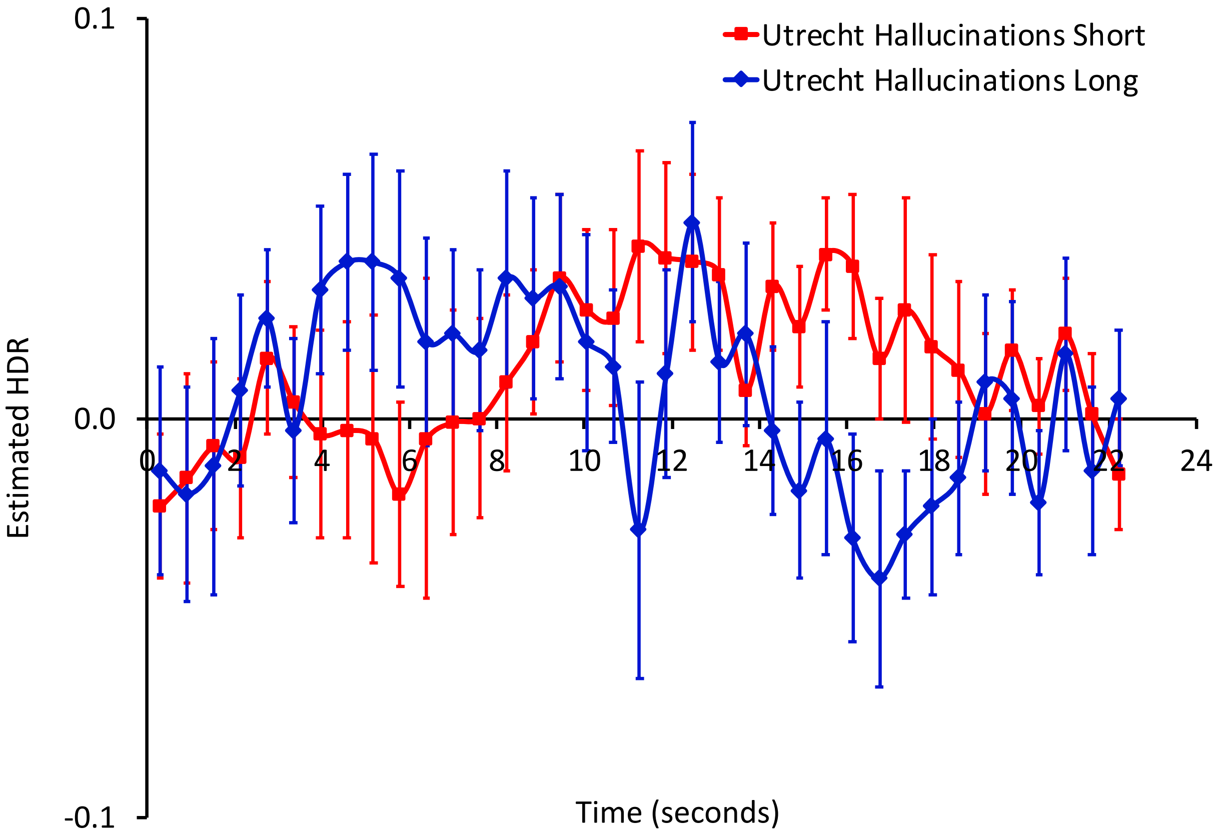

**Supplementary Figure S5:** *Melbourne only hallucinations Component 1. A (top): dominant 20% of component loadings for Component 1, from the Melbourne patient hallucinations (S/M/L) analysis. MNI Z-axis coordinates are displayed. Images are displayed in neurological convention (left is left). Red/yellow = positive loadings, positive threshold = 0.13, max = 0.21. B (bottom): mean finite impulse response (FIR)-based predictor weights plotted as a function of post-stimulus time bin (TR = 1150ms) and condition (averaged over participants, error bars are standard errors).*

A

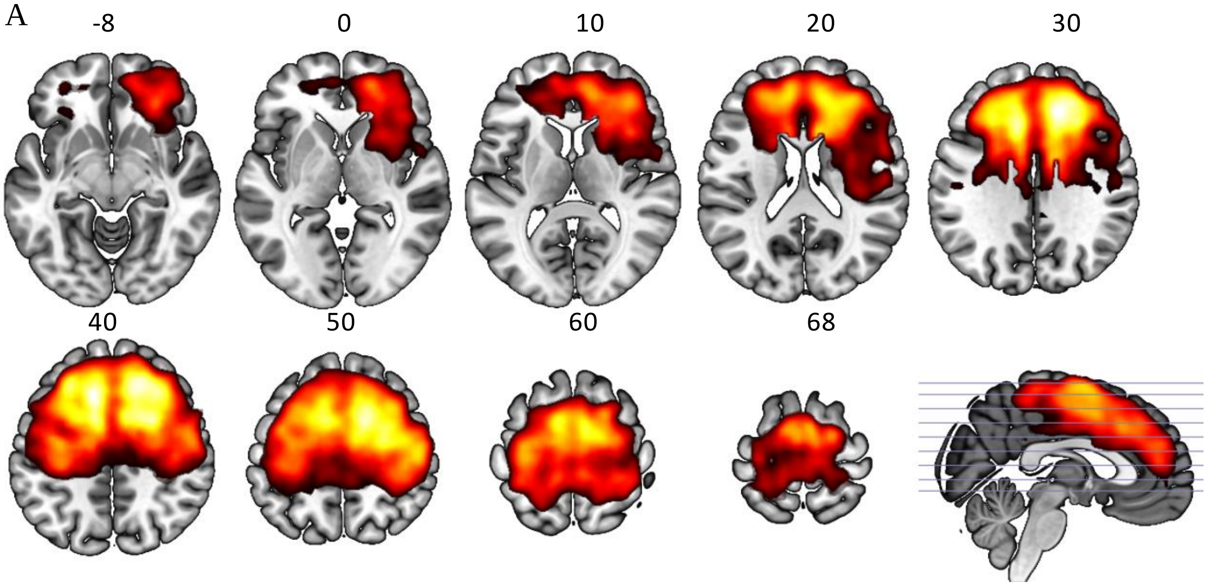


B

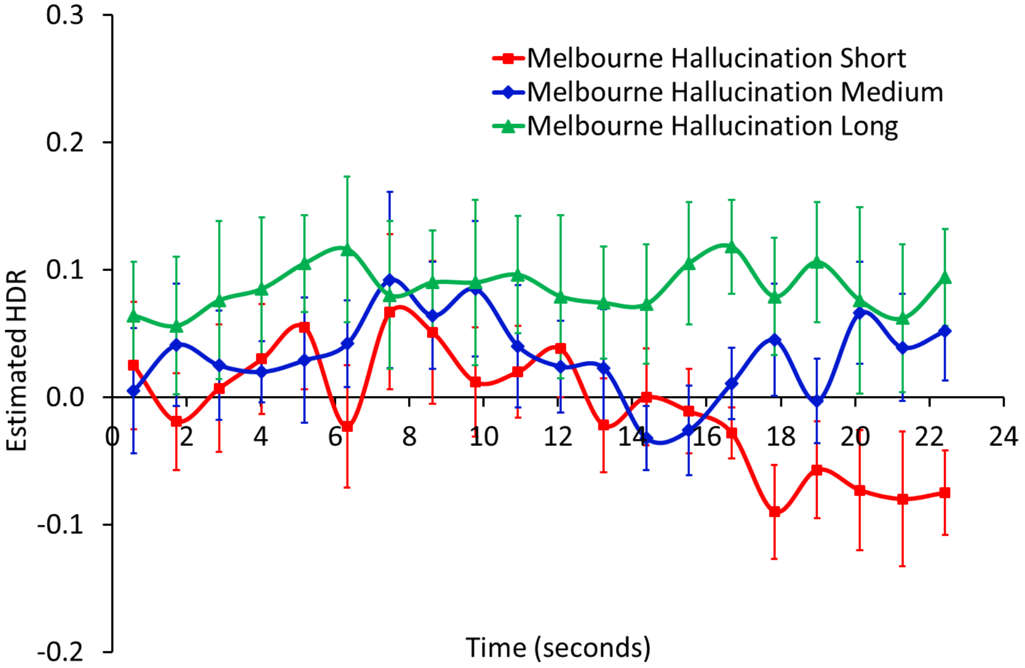

**Supplementary Figure S6:** *Melbourne alone hallucinations Component 2. A (top): dominant 20% of component loadings for Component 2, proposed FVF network, from the Melbourne alone hallucinations (S/M/L) analysis. MNI Z-axis coordinates are displayed. Images are displayed in neurological convention (left is left). Blue/green = negative loadings, negative threshold = -0.13, min = -0.21. B (bottom): mean finite impulse response (FIR)-based predictor weights plotted as a function of post-stimulus time (TR = 1150ms) and condition (averaged over participants, error bars are standard errors).*

*A*

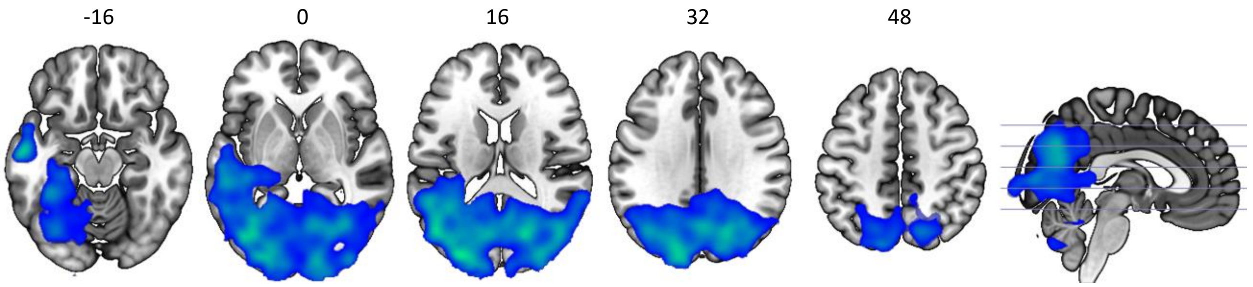


B

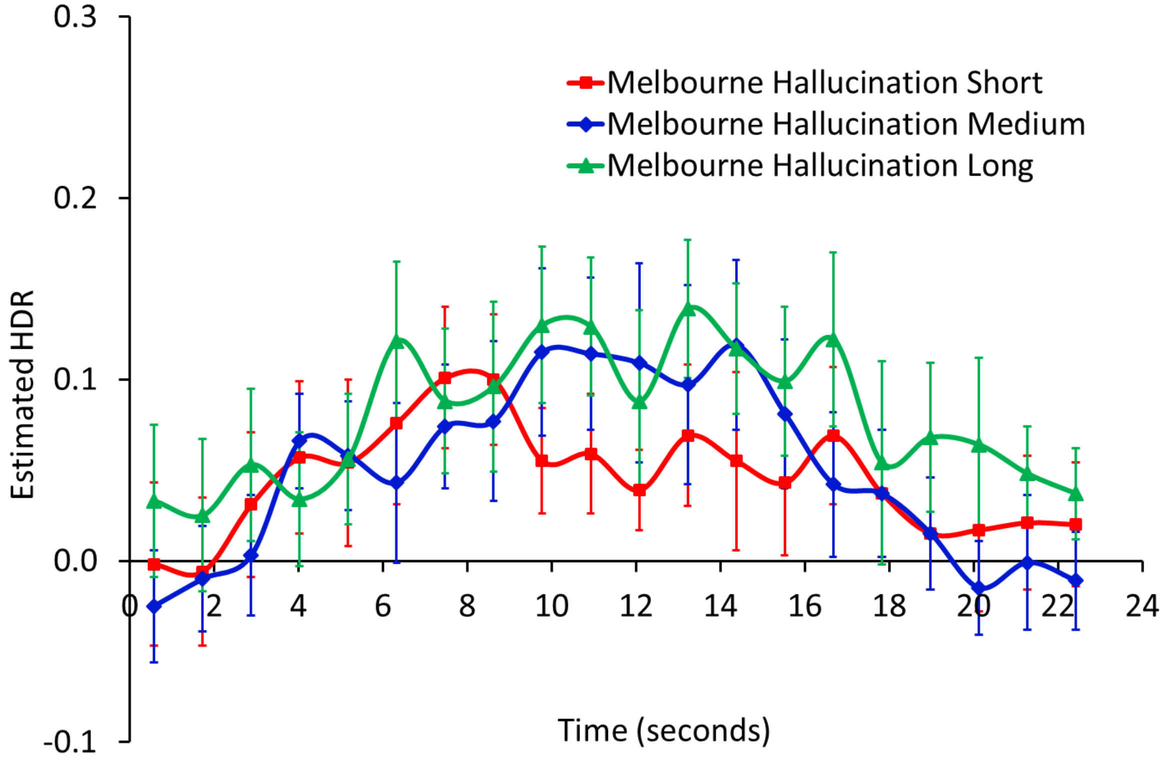

**Supplementary Figure S7:** *Melbourne alone hallucinations Component 3. A (top): dominant 20% of component loadings for Component 3, from the Melbourne patient internal (S/M/L) analysis. MNI Z-axis coordinates are displayed. Images are displayed in neurological convention (left is left). Red/yellow = positive loadings, positive threshold = 0.10, max = 0.25. B (bottom): mean finite impulse response (FIR)-based predictor weights plotted as a function of post-stimulus time bin (TR = 1150ms) and condition (averaged over participants, error bars are standard errors).*

A

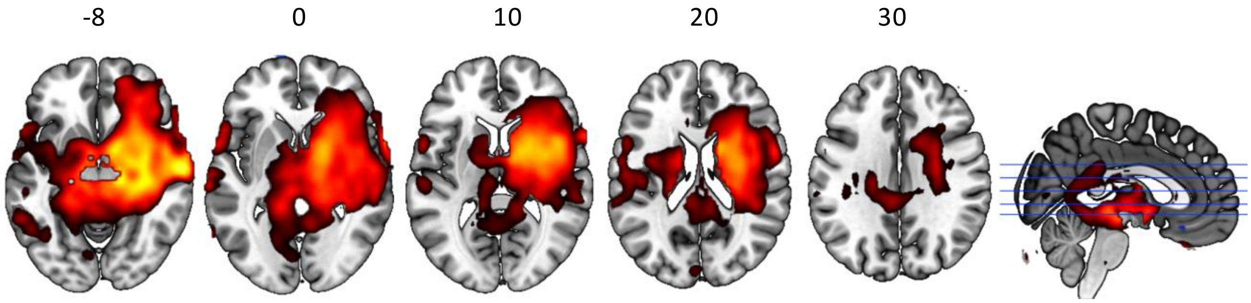


B

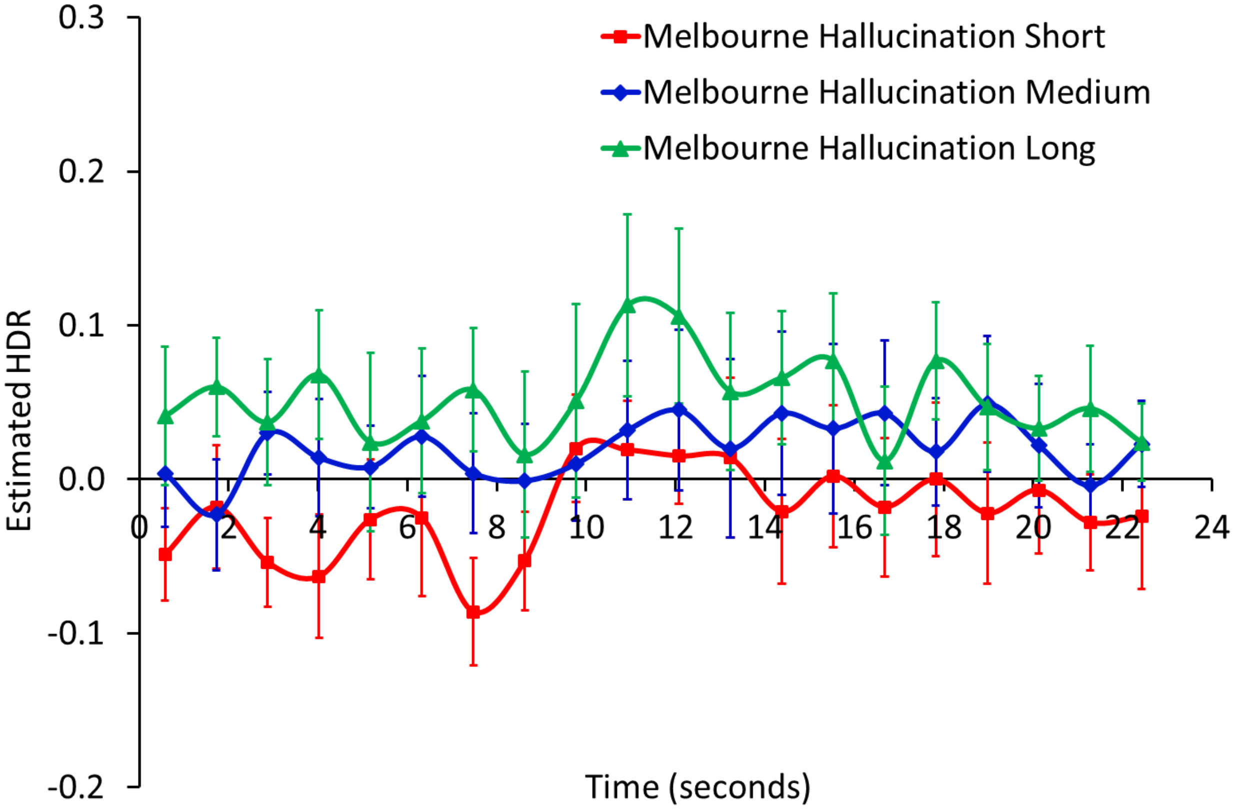

**Supplementary Figure S8:** *Melbourne alone hallucinations Component 4. A (top): dominant 20% of component loadings for Component 4, from the Melbourne patient internal (S/M/L) analysis. MNI Z-axis coordinates are displayed. Images are displayed in neurological convention (left is left). Blue/green = negative loadings, negative threshold = -0.10, min = -0.24. B (bottom): mean finite impulse response (FIR)-based predictor weights plotted as a function of post-stimulus time bin (TR = 1150ms) and condition (averaged over participants, error bars are standard errors).*

A

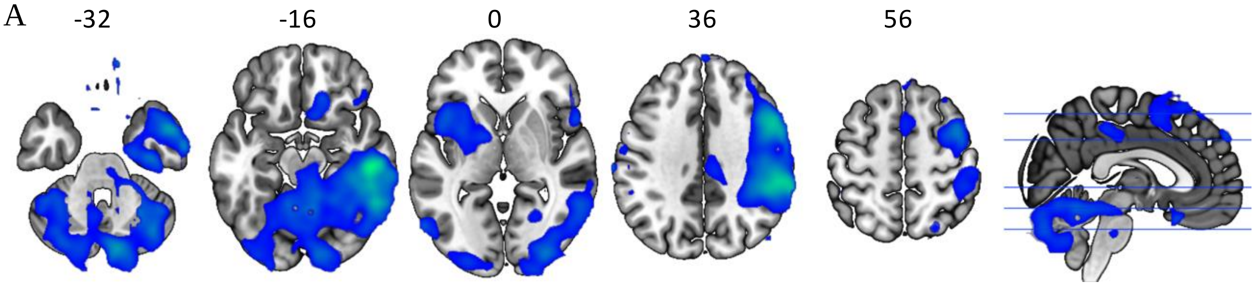


B

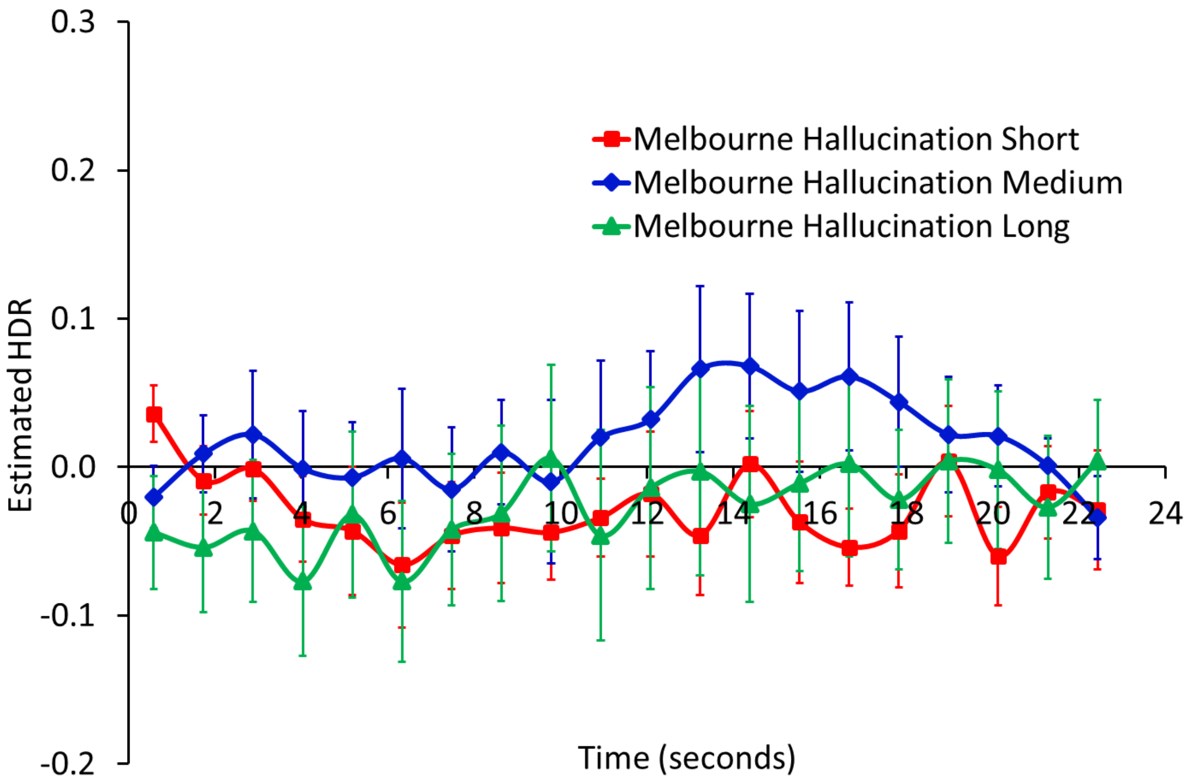

**Supplementary Figure S9:** *Utrecht alone hallucinations (S/L) Component 1. A (top): dominant 20% of component loadings for Component 1, proposed two-handed response (2RESP) network, from the Utrecht patient internal (S/L) analysis. MNI Z-axis coordinates are displayed. Images are displayed in neurological convention (left is left). Red/yellow = positive loadings, positive threshold = 0.10, max = 0.32. B (bottom): mean finite impulse response (FIR)-based predictor weights plotted as a function of post-stimulus time (TR = 610ms) and condition (averaged over participants, error bars are standard errors).*

A

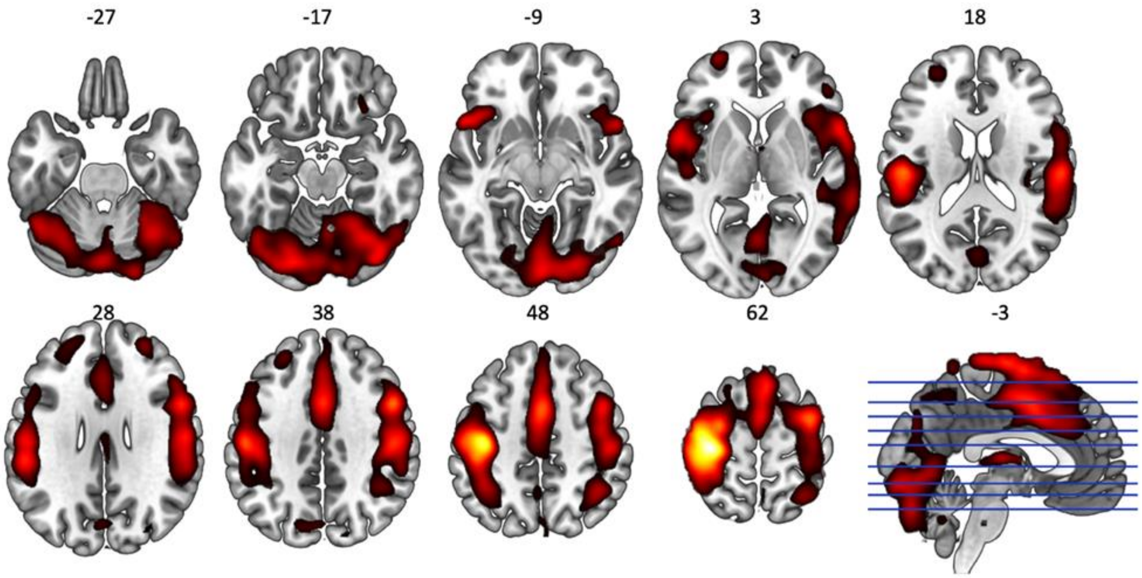


B

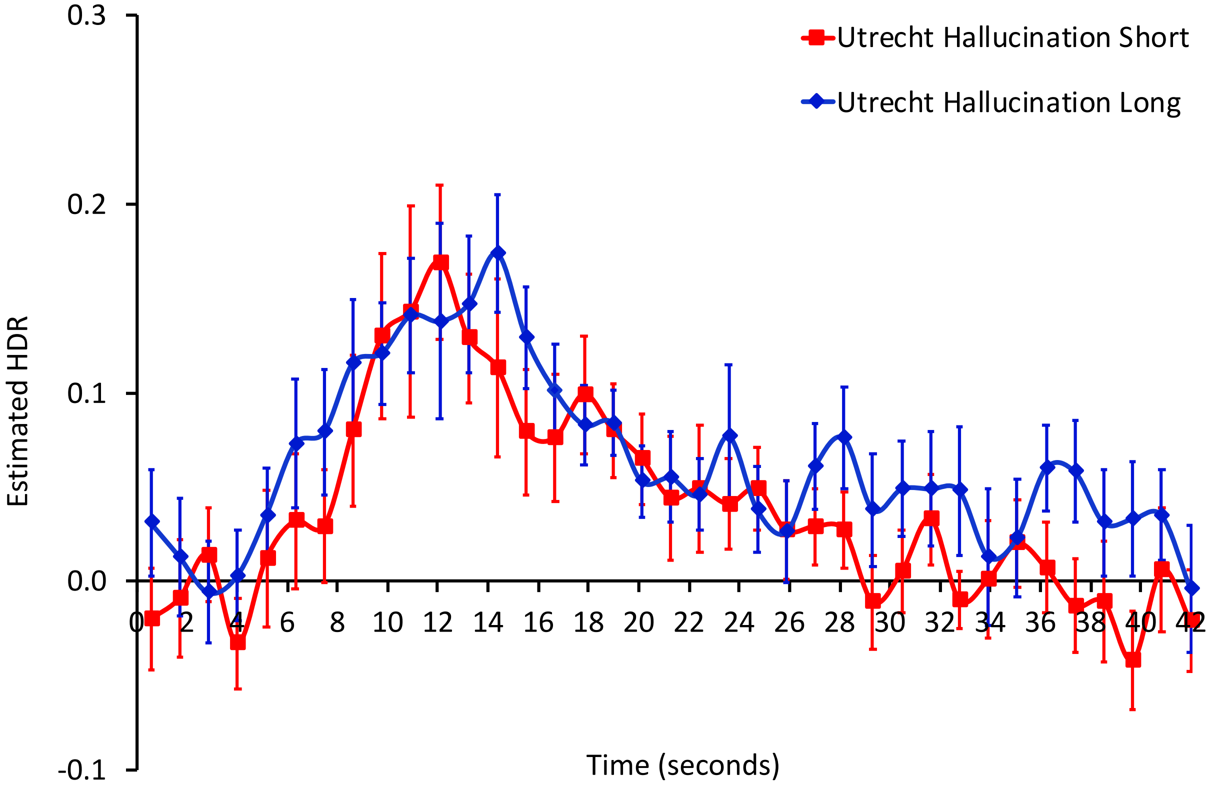

**Supplementary Figure S10:** *Utrecht alone hallucinations (S/L) Component 2. A (top): dominant 20% of component loadings for Component 2, from the Utrecht alone patient hallucinations (Short/Long) analysis. MNI Z-axis coordinates are displayed. Images are displayed in neurological convention (left is left). Red/yellow = positive loadings, positive threshold = 0.09, max 0.17. B (bottom): mean finite impulse response (FIR)-based predictor weights plotted as a function of post-stimulus time (TR = 610ms) and condition (averaged over participants, error bars are standard errors).*

A

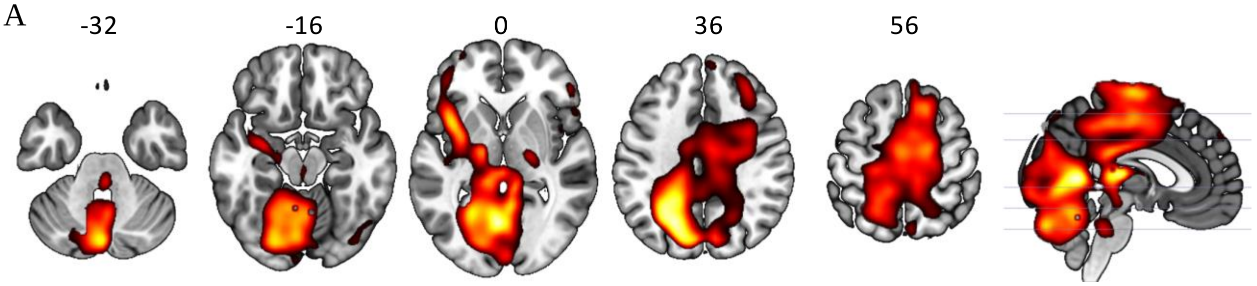


B

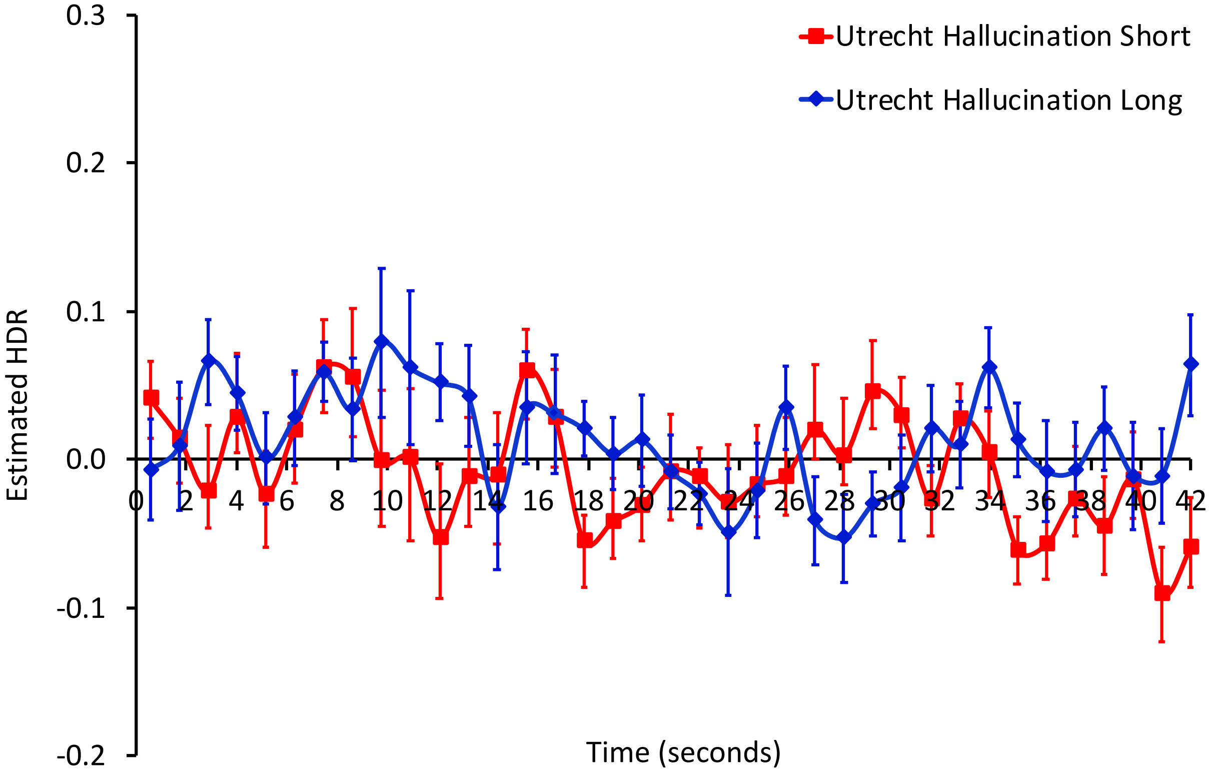
